# Measuring the selective packaging of RNA molecules by viral coat proteins in cells

**DOI:** 10.1101/2025.04.14.648219

**Authors:** Amineh Rastandeh, Nino Makasarashvili, Herman K. Dhaliwal, Sundharraman Subramanian, Daniel A. Villarreal, Sherry Baker, Elmer I. Gamez, Kristin N. Parent, Rees F. Garmann

**Affiliations:** Department of Chemistry and Biochemistry, San Diego State University, San Diego, CA 92182; Department of Biochemistry and Molecular Biology, Michigan State University, East Lansing, MI 48824; Viral Information Institute, San Diego State University, San Diego, CA 92182

**Keywords:** plus-strand RNA virus, capsid assembly, RNA packaging, packaging signals

## Abstract

Some RNA viruses package their genomes with extraordinary selectivity, assembling protein capsids around their own viral RNA while excluding nearly all host RNA. How the assembling proteins distinguish viral RNA from host RNA is not fully understood, but RNA structure is thought to play a key role. To test this idea, we perform in-cellulo packaging experiments using bacteriophage MS2 coat proteins and a variety of RNA molecules in *E. coli*. In each experiment, plasmid-derived RNA molecules with a specified sequence compete against the cellular transcriptome for packaging by plasmid-derived coat proteins. Following this competition, we quantify the total amount and relative composition of the packaged RNA using electron microscopy, interferometric scattering microscopy, and high-throughput sequencing. By systematically varying the input RNA sequence and measuring changes in packaging outcomes, we are able to directly test competing models of selective packaging. Our results rule out a longstanding model in which selective packaging requires the well-known TR stem-loop, and instead support more recent models in which selectivity emerges from the collective interactions of multiple coat proteins and multiple stem-loops distributed across the RNA molecule. These findings establish a framework for understanding selective packaging in a range of natural viruses and virus-like particles, and lay the groundwork for engineering synthetic systems that package specific RNA cargoes.

**Significance Statement:** Bacteriophage MS2 packages its RNA genome into protective protein shells called capsids while excluding nearly all host-cell RNA. Engineering synthetic capsids with similar selectivity could enable a broad range of RNA-based technologies, including CRISPR gene editing systems, mRNA vaccines, and other emerging RNA-based therapeutics. Our study shows that selective packaging in MS2 is not dictated by a single, high-affinity RNA-protein interaction but instead emerges from the collective interactions of multiple coat proteins and an ensemble of stem-loops distributed across the RNA molecule. By establishing these collective interactions as the basis of selectivity, our findings provide a foundation for engineering synthetic capsids capable of selectively packaging target RNAs for next-generation RNA-based technologies.

## Introduction

Many plus-strand RNA viruses package their RNA genomes into protective protein shells called capsids (1). For many of these viruses, genome packaging and capsid formation occur together as a concerted process in which the viral coat proteins assemble around the viral RNA to form the capsid (2). This process takes place in the cytoplasm of the host cell, a crowded environment filled with host RNA molecules (3). Despite the abundance of host RNA, some viruses manage to package their own RNA with greater than 99% fidelity, meaning that less than 1% of the encapsidated RNA content belongs to the host (4–6). How the assembling proteins distinguish viral RNA from host RNA is not completely understood, but the viral RNA molecules are thought to adopt specific structures termed “packaging signals” (7) that facilitate selection.

For over 50 years, bacteriophage MS2 has served as a model system for studying RNA packaging (8–10), but its packaging signal remains an open question (**Fig. 1A**). Early studies (11, 12) proposed that MS2 packaging is driven by a specific, high-affinity interaction between the MS2 coat protein and a single stem-loop in the MS2 RNA, known as the “TR-loop” (13). However, more recent structural (14, 15) and biochemical (16) studies suggest that additional regions of the RNA molecule—potentially a dozen or more stem-loops distributed across the primary sequence—might also play an important role (**Fig. 1B**). Still other studies raise an alternative hypothesis: that packaging does not rely on locally folded stem-loops but instead depends on “global” (17) properties of the RNA, such as the total length and electrostatic charge (18) or the overall physical size and shape (19). Distinguishing between these competing hypotheses, and establishing a working model of the MS2 packaging signal, requires direct measurements of packaging selectivity carried out systematically across a variety of RNA molecules with different sequences and structures. Surprisingly, despite decades of MS2 packaging research, such experiments have not been reported.

**Fig. 1.**
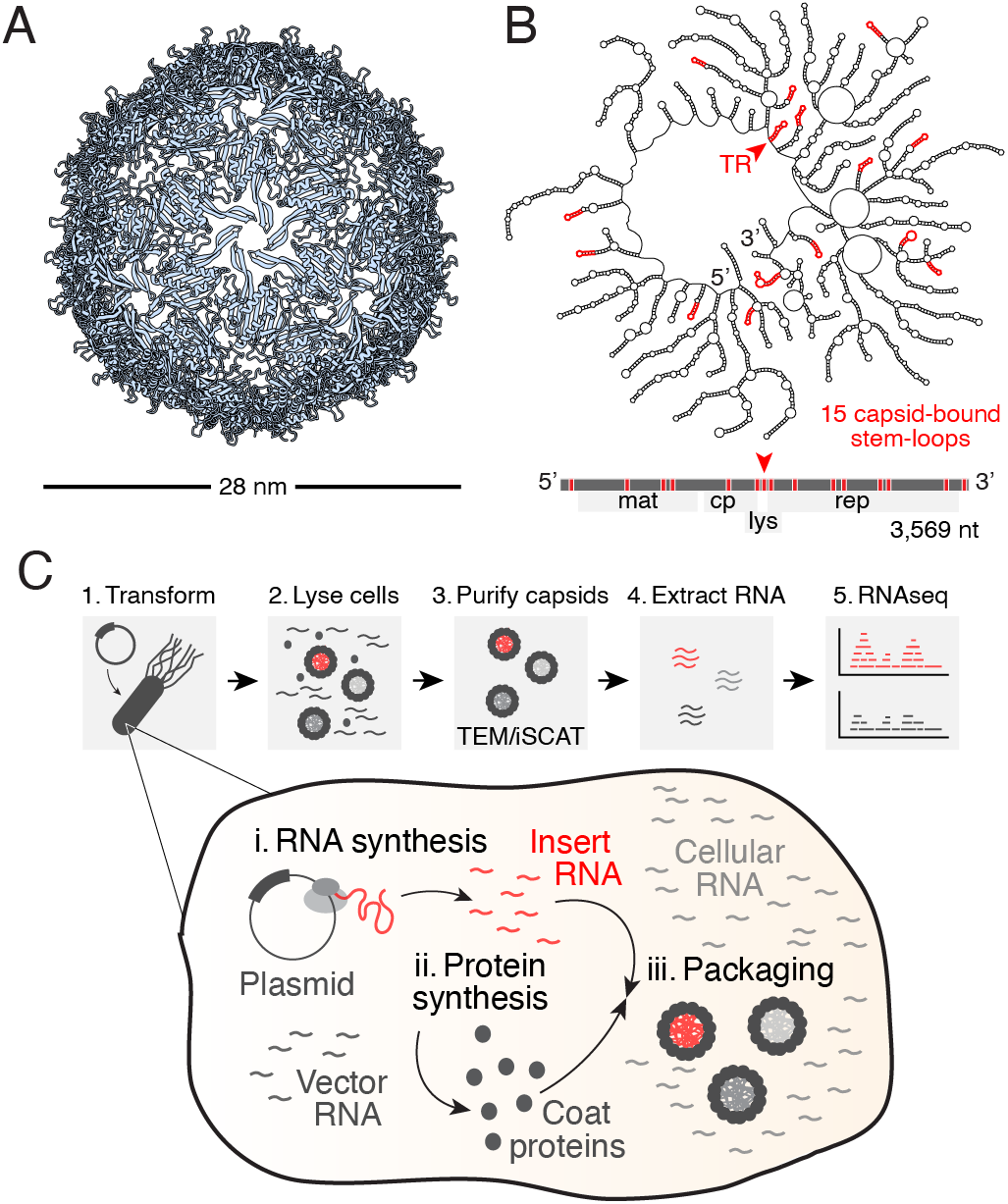
Overview of the system and the experiment. **A**. A structural model of the MS2 capsid contains 180 copies of the coat protein arranged as a 28-nm icosahedral shell (21). **B**. A secondary structure model of the packaged MS2 genome contains 15 stem-loop structures (shown in red) that bind tightly to the interior surface of the capsid (15). One of these stem-loops is the famous TR loop (13) (shown by a red arrowhead). A map of the primary structure shows the positions of these stem-loops relative to the viral genes. **C**. A schematic of the in-cellulo packaging experiment outlines the key steps of our protocol: **1**. We transform *E. coli* with a plasmid whose insert contains the MS2 coat protein gene flanked on either side by variable untranslated sequences. **Inset:** Once transformed, the plasmid insert is transcribed into RNA (**i**), which is then translated into coat protein (**ii**). When the cellular concentration of coat protein becomes sufficiently high, capsids assemble and package some of the available pool of RNA (**iii**). This pool contains a mixture of insert transcripts, plasmid vector-derived transcripts, and cellular transcripts, all of which compete for packaging by the assembling coat proteins. **2**. We lyse the cells after 24 h to release the assembled capsids, and then **3**. purify the capsids from unpackaged RNA and other cellular debris using nuclease digestion followed by ultracentrifugation. Once purified, we can infer the amount of RNA packaged per particle using a combination of electron microscopy (TEM) and interferometric scattering microscopy (iSCAT). Finally, **4**. we extract the packaged RNA from the capsids, and **5**. determine its identity using short-read high-throughput RNA sequencing (RNAseq).

To address this problem, we develop a simple in-cellulo packaging experiment that enables direct measurements of RNA packaging by MS2 coat proteins in *E. coli* cells (**Fig. 1C**). Our approach is inspired by recent in-planta experiments on the packaging of RNA by satellite tobacco necrosis virus (20). In our experiment, plasmid-derived RNA molecules expressed in *E. coli* compete with the cellular transcriptome for packaging by plasmid-derived MS2 coat proteins. Following this packaging competition, we use electron microscopy, interferometric scattering microscopy (iSCAT), and high-throughput RNA sequencing to determine the amount and identity of the packaged RNA, providing a quantitative measure of selectivity (**Fig. 1C**). By systematically varying the properties of the input RNA molecules—including length, physical size, and the presence or absence of specific stem-loops—and then measuring packaging outcomes, we investigate how selectivity depends on the RNA structure. We use our results to critically evaluate current hypotheses of the MS2 packaging signal, with the goal of establishing design rules for building synthetic capsids with high selectivity.

## Results and discussion

### MS2 coat protein alone can selectively package MS2 RNA

While natural MS2 packages its genome into capsids containing 178 coat proteins and a single maturase protein (15), our experiments focus on RNA packaging into capsids composed solely of coat protein. By omitting the maturase protein, a specialized component unique to certain RNA bacteriophages, we aim to uncover general principles of RNA packaging that apply to a broader range of viruses and virus-like particles whose capsids consist of only coat protein (22–27).

To test whether MS2 coat-protein-only capsids can achieve high selectivity, we performed an initial experiment involving a modified version of the native MS2 genome. We constructed a pET expression plasmid, pMS2′, containing an insert that encodes the full 3,569-nt MS2 genome, modified with point mutations introducing early stop codons in all viral genes except the coat protein gene (see **Methods**; **Fig. 2A, top**; and **SI Appendix, Fig. S1** and **Dataset S1**). When expressed in the cell, pMS2′ transcripts maintain nearly the same sequence as natural MS2 RNA, but they produce only coat proteins upon translation.

**Fig. 2.**
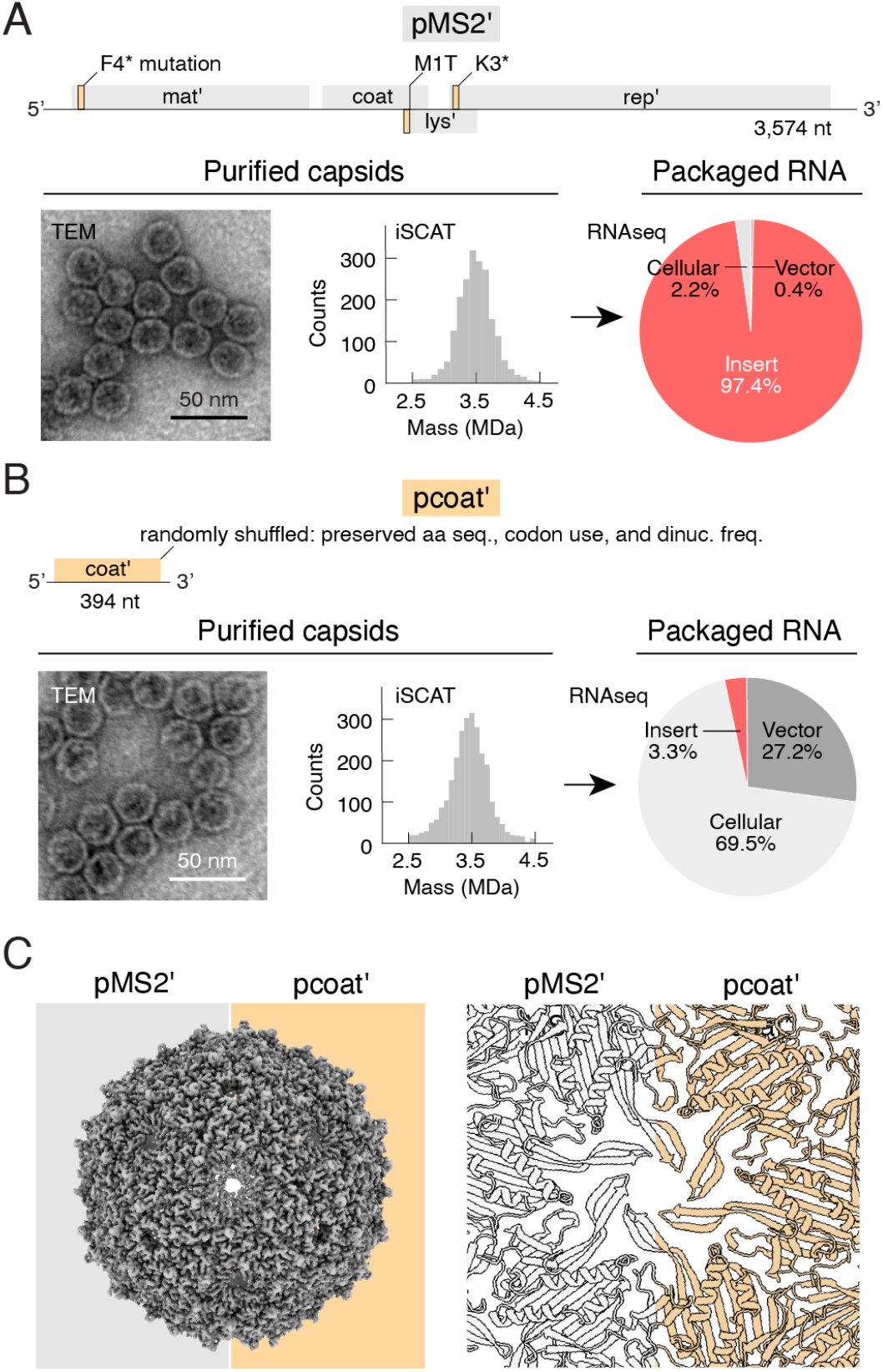
Varying the insert sequence yields similar capsids with different RNA contents. **A**. The pMS2′ insert contains the full MS2 genome (**top**), modified with point mutations to prevent the production of viral proteins other than the coat protein (**SI Appendix, Fig. S1**). Leaky expression of pMS2′ transcripts produced well-formed 28-nm capsids, as observed by negative-stain TEM (**bottom-left**), with an average mass of 3.5 MDa per particle, determined by iSCAT (**bottom-middle**). Approximately 97% (±1% across duplicate experiments) of sequencing reads from the packaged RNA aligned to the pMS2′ insert sequence (**bottom-right**). **B**. The pcoat′ insert contains only the MS2 coat protein gene (**top**), modified by random codon swaps to scramble the RNA sequence while preserving the amino acid sequence, codon usage, and dinucleotide frequency (**SI Appendix, Fig. S5**). As with pMS2′, leaky expression of pcoat′ transcripts produced well-formed 28-nm capsids (**bottom-left**) with an average mass of 3.5 MDa (**bottom-middle**). However, only 3% (±1% across duplicate experiments) of sequencing reads aligned to the insert sequence, with 27% aligning to the plasmid vector and 70% to the *E. coli* (cellular) genome (**bottom-right**). **C**. Cryo-electron microscopy and single-particle reconstruction showed that the majority of particles produced by both inserts adopt identical capsid structures. The electron density maps of each capsid overlap (**left**), and their molecular models are indistinguishable (**right**).

Leaky expression of pMS2′ insert transcripts—without IPTG induction—results in the formation of well-ordered, RNA-containing capsids. Coat proteins translated from the insert transcripts assemble into monodisperse spherical capsids with size and curvature consistent with natural MS2, as observed by negative-stain transmission electron microscopy (TEM) (**Fig. 2A, bottom-left**). These capsids package nucleic acids, as evidenced by a dominant peak at 260 nm in their UV-absorbance spectra (**SI Appendix, Fig. S2**). Furthermore, the packaged nucleic acid is predominantly RNA, as confirmed by its complete digestion with RNase A following extraction from the capsid using phenol:chloroform (**SI Appendix, Fig. S2**).

We quantified the total mass of RNA packaged per particle using interferometric scattering microscopy (iSCAT) (28) (see **Methods** and **SI Appendix, Fig. S3**). The mass distribution of pMS2′ particles, determined from 2,000 single-particle measurements, reveals a clear peak at 3.5 MDa (**Fig. 2A, bottom-middle**), closely matching the expected mass of 3.6 MDa for natural MS2. Given the structural similarity between pMS2′ particles and natural MS2 (**Fig. 2A, bottom-left**), we infer that both particle types contain comparable protein mass and, consequently, similar RNA mass. Assuming pMS2′ particles contain 2.5 MDa of protein—equivalent to natural MS2—we estimate that these particles package approximately 1 MDa, or 3.3 knt, of RNA. This value is within 10% of the 3.6 knt packaged by natural MS2.

To determine the identity of the packaged RNA, we extracted it from the capsids and analyzed it using high-throughput short-read sequencing (see **Methods** and **SI Appendix, Fig. S4**). To interpret the sequencing data, we first aligned the reads to the plasmid sequence, differentiating between those aligning to the insert portion and the vector portion, which contains genes for kanamycin resistance, the LacI repressor, and other regulatory elements (**SI Appendix, Fig. S1**). Reads that did not align to the plasmid were subsequently aligned to the *E. coli* genome (Genbank CP053602.1; **SI Appendix, Dataset S2**). Using this approach, we found that 97.4% of the total reads aligned to the pMS2′ insert, 0.4% to the vector, 2.2% to the *E. coli* genome, and 0.02% failed to align (**Fig. 2A, bottom-right**). These values represent the averages of two independent biological experiments, which agree to within 0.4% (**SI Appendix, Table S1**).

We define the ‘packaging fraction’ of an insert as the percentage of packaged sequencing reads that align to it (see **Methods**). For the pMS2′ insert, the packaging fraction is roughly 97%, indicating that nearly all pMS2′ particles contain an insert transcript. However, because our sequencing measurements average across many particles, it remains unclear whether the 3% minority fraction of vector and cellular transcripts is packaged at high density in a small subset of particles or co-packaged with insert transcripts at low density across many particles.

Next, we compared the packaging fraction of the pMS2′ insert to the prevalence of insert transcripts in the cell, which we measured by sequencing the total cellular RNA. To prevent the accumulation of packaged RNA in the cell, we introduced an additional mutation into the pMS2′ insert sequence, creating an early stop codon in the coat gene to abolish packaging (see **SI Appendix, Fig. S1**). Sequencing the total cellular RNA in the absence of packaging revealed that pMS2′ insert transcripts constitute less than 2% of the total RNA in the cell (**SI Appendix, Fig. S4**). These results suggest that the 97% packaging fraction observed for the pMS2′ insert is not merely due to transcript abundance, but instead reflects selective packaging of the insert transcripts over cellular RNA.

To test whether the selective packaging of insert transcripts depends on their sequence, we repeated the packaging experiment using a different insert. We constructed a new expression plasmid, pcoat′, containing a minimal insert that encodes only the 394-nt coat protein gene (**Fig. 2B, top**). To distinguish the pcoat′ insert from the pMS2′ insert, we randomly shuffled its nucleotide sequence while preserving codon usage and dinucleotide frequency (29, 30) (see **Methods**, and **SI Appendix, Fig. S5**). This design ensures consistent protein production levels and minimizes complications arising from atypical dinucleotide distributions (31). When expressed in the cell, pcoat′ transcripts display limited RNA sequence similarity to the coat gene portion of pMS2′ transcripts. However, translation of these transcripts produces identical coat proteins, which assemble into capsids and package RNA.

The assembled pcoat′ capsids are structurally similar to pMS2′ capsids but primarily package cellular RNA molecules. TEM shows that pcoat′ and pMS2′ particles have comparable sizes and shapes (**Figs. 2A and 2B, bottom-left**), and iSCAT indicate equivalent masses (**Figs. 2A and 2B, bottom-middle**), suggesting that both types of particles have similar capsid structures and similar amounts of RNA. However, the RNA composition differs considerably. For pcoat′ particles, only 3.3% of the packaged reads align to the insert, while 27.2% align to the vector, 69.9% to the *E. coli* genome, and 0.06% fail to align (**Fig. 2B, bottom-right**). Thus, the packaging fraction of the pcoat′ insert is only 3%. From this value, we infer that at most one in four pcoat′ particles contains a complete insert transcript (see **SI Appendix, Supporting text**).

A detailed comparison of pMS2′ and pcoat′ particles using single-particle cryo-electron microscopy approaches and image reconstructions reveals highly similar capsid structures, with minor differences. For both constructs, the majority of particles—99.3% of pMS2′ particles and 92.2% of pcoat′ particles—exhibit the canonical T=3 capsid structure. The reconstructed electron density maps of T=3 capsids for pMS2′ and pcoat′ are essentially identical (**Fig. 2C, left**), and molecular models of their protein structures overlap with a root mean square deviation of 0.045 Å (**Fig. 2C, right**). Minority populations of non-canonical structures were also observed for both constructs: 0.7% of pMS2′ capsids and 2% of pcoat′ capsids adopt a recently reported D5-symmetric prolate structure (32), and 5.8% of pcoat′ particles exhibit a previously unreported oblate structure (see **SI Appendix, Figs. S6 and S7**). Thus, while pcoat′ particles appear slightly more heterogeneous, the predominant species for both constructs remains the canonical T=3 structure.

Together, these results demonstrate that MS2 coat-protein-only capsids adopt well-ordered structures and package RNA. While the total amount of packaged RNA appears to be independent of the insert sequence, the RNA composition is strongly influenced by the insert sequence. To identify the key properties of RNA molecules that affect selectivity, we performed packaging experiments using a series of inserts with varying lengths and sequences, as described below.

### RNA features that impact selectivity Length

The most apparent difference between the pMS2′ and pcoat′ inserts is their length. At 3,574 nts, pMS2′ transcripts are longer than all vector-derived transcripts and most *E. coli* transcripts in the cell, including the highly abundant rRNA transcripts (3) (**SI Appendix, Fig. S8**). By contrast, pcoat′ transcripts, at 394 nts, are considerably shorter than these other RNA molecules.

Could length alone account for the observed difference in packaging selectivity? To address this question, we constructed a series of additional insert sequences ranging in length from 394 to 3,570 nts (**Fig. 3A**). Each insert includes the MS2 coat protein gene followed by varying lengths of random non-coding sequence. Unlike the pcoat′ insert, which contains a shuffled version of the coat gene, these inserts use the unshuffled coat gene to allow direct comparisons with subsequent experiments.

**Fig. 3.**
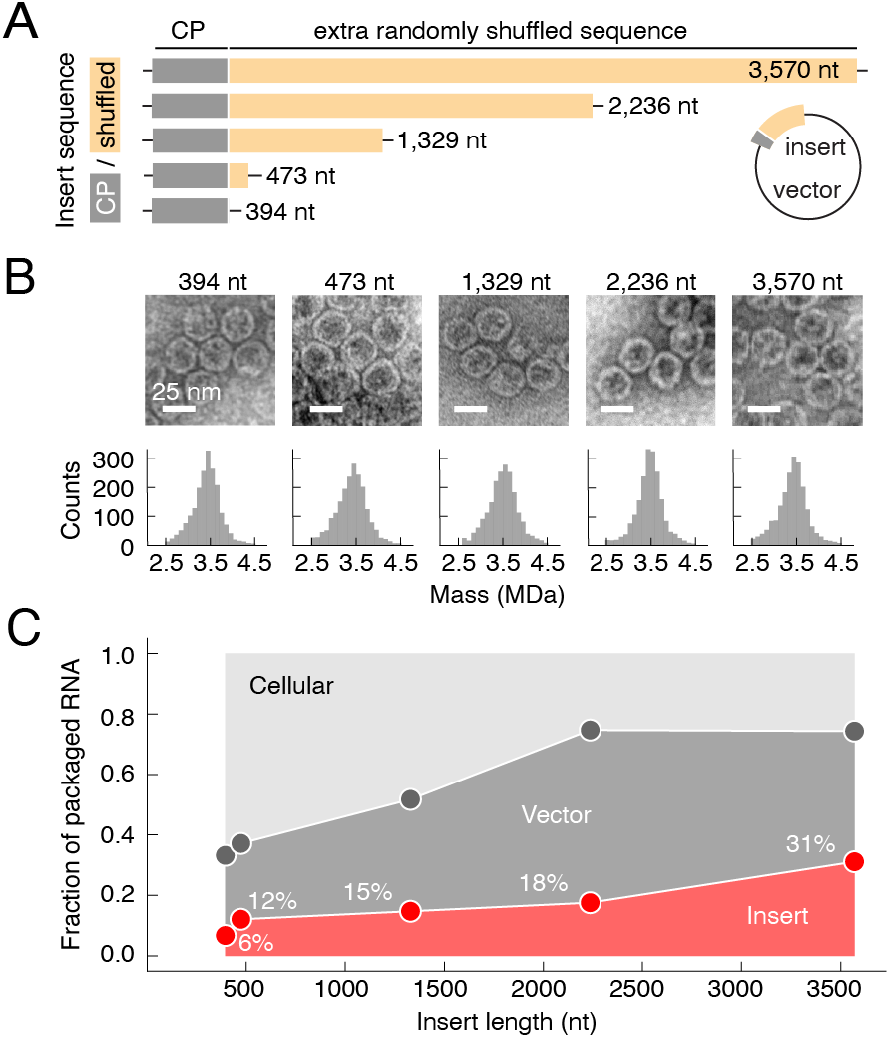
Selectivity depends weakly on RNA length. **A**. Inserts of varying lengths were constructed by appending random dinucleotide-preserving shuffled sequences downstream of the coat protein gene. **B**. For all lengths tested, TEM confirmed the formation of well-formed 28-nm capsids (**top**), while iSCAT measurements showed an average particle mass of 3.5 MDa (**bottom**). **C**. The fraction of packaged RNA sequencing reads aligning to the insert, vector, or host genome is plotted as a function of insert length. The results show that the insert packaging fraction increases monotonically with length. However, these fractions remain well below the 97% observed for pMS2′, indicating that RNA length alone is insufficient to achieve high selectivity.

For each insert tested, the assembled particles displayed the expected 28-nm spherical morphology by TEM and a consistent 3.5-MDa mass by iSCAT (**Fig. 3B**), suggesting that all particles package a similar amount of RNA. However, the RNA composition—specifically, the packaging fraction of the insert transcripts—varied with insert length. We observed a monotonic increase in packaging fraction with increasing length, from 6% for the shortest 394-nt insert to 31% for the longest 3,570-nt insert (**Fig. 3C**). These findings suggest that RNA length does influence packaging selectivity. However, length alone cannot account for the high packaging fractions observed with the pMS2′ insert (97%, **Fig. 2A**). In the following experiments, in addition to length, we systematically vary the RNA sequence and structure to determine key features that give rise to high selectivity.

### Packaging signals vs. physical size

Competing hypotheses posit distinct roles for RNA sequence and structure in packaging selectivity. One prominent hypothesis proposes that RNA packaging signals—locally folded stem-loop structures that bind tightly to MS2 coat proteins—are critical for selection (33). According to this model, strong binding between packaging signals and coat proteins promotes capsid assembly, making RNA molecules with these signals more likely to be packaged (34). An alternative hypothesis emphasizes the importance of global properties, such as the overall physical size and shape of the RNA molecule (35). This model suggests that RNA sequences adopting compact structures fit more efficiently within the limited interior volume of the capsid, leading to preferential packaging of compact molecules over extended ones (35).

To differentiate between these hypotheses, we designed a set of insert sequences with varying local and global structures (**Fig. 4A**). The design process began with a base sequence, derived as a circular permutation of the pMS2′ insert, starting at the coat gene. From this base sequence, we systematically shuffled nucleotides within specific regions—namely, those proposed to contain packaging signal stem-loops and the regions between them—while preserving the overall dinucleotide frequency (**SI Appendix, Fig. S9**). This strategy yielded five distinct shuffle types, as shown in **Fig. 4A, left**:

**Fig. 4.**
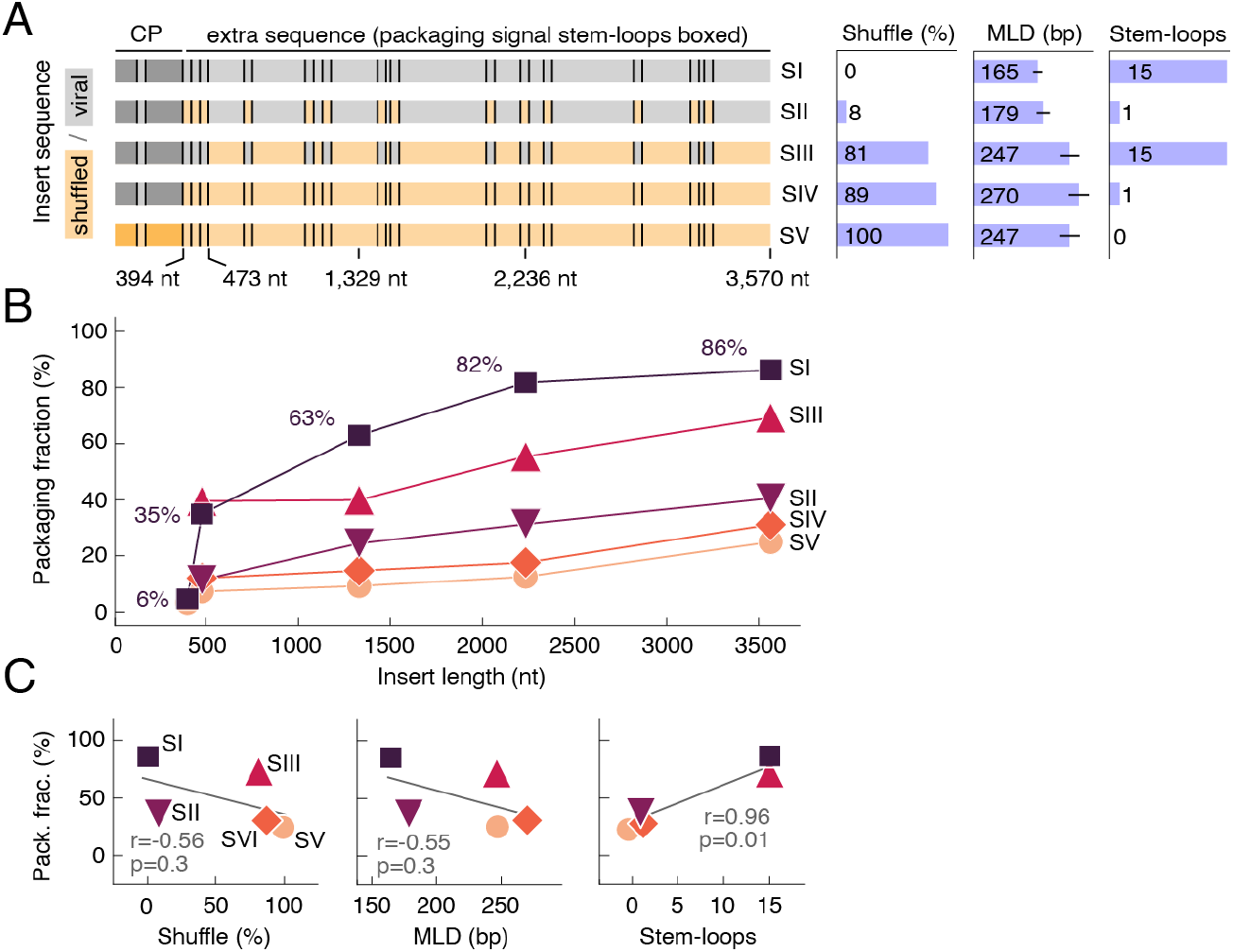
Selectivity depends strongly on the presence of special stem-loops. **A**. **Left:** The insert sequence was systematically varied by appending portions of the MS2 sequence downstream of the coat gene and shuffling nucleotides within specific regions, yielding five shuffle types, SI–SV. For detailed descriptions of these shuffle types, see the **main text** and **SI Appendix, Dataset S1. Right:** Bar plots display the fraction of shuffled nucleotides, the maximum ladder distance (MLD), and the number of packaging signal stem-loops for each shuffle type. MLD, a theoretical measure of RNA molecule size derived from thermodynamic folding models, is displayed as the mean (blue bars) and standard deviation (black bars) calculated across 1,000 predicted equilibrium secondary structures (see **Methods** and **SI Appendix, Fig. S11**). **B**. packaging fractions of each insert are shown as a function of length. Shuffle types are indicated by distinct symbols, connected by straight lines to guide the eye. **C**. The packaging fraction is shown as a function of the fraction of shuffled nucleotides (**left**), MLD (**middle**), and the number of stem-loops (**right**) for the full-length transcripts of each shuffle type. Best-fit regression lines (gray) are included, along with Pearson correlation coefficients (r) and p-values (p).

- **SI (Shuffle type I)**: The unshuffled base sequence.
- **SII**: Shuffled in regions corresponding to 14 proposed packaging signal stem-loops.
- **SIII**: Shuffled in the regions between the 14 stem-loops.
- **SIV**: Completely shuffled downstream of the coat gene (as in **Fig. 3)**.
- **SV**: Fully shuffled across the entire insert, including the coat gene, in which the amino acid sequence and overall condon frequency are preserved (as in **Fig. 2B**).

These shuffle types differ in their fraction of shuffled nucleotides, the number of packaging signal stem-loops, and their physical sizes (**Fig. 4A, right**). Although directly measuring the physical size of an RNA molecule in solution is challenging, theoretical estimates can be derived from RNA folding models (36). Specifically, the maximum ladder distance (MLD) estimates the size of an RNA molecule by quantifying the extendedness of its predicted secondary structures (19). Previous studies have demonstrated that MLD values correlate well with experimental measurements of RNA hydrodynamic radii in solution (37). Using this approach, we calculated the MLD for each shuffle type and found that SI and SII exhibit significantly smaller MLDs—and thus more compact structures—than SIII, SIV, and SV (**Fig. 4A, right, “MLD (bp)”**; and **SI Appendix, Figs. S10 and S11**). The compact structure of SI aligns with previous findings that natural MS2 RNA is exceptionally compact (38).

By varying the length and shuffle type of the inserts, we observed a broad range of packaging fractions and identified some general trends. As expected, packaging fractions increased with insert length within each shuffle type (**Fig. 4B**), consistent with our earlier observation that longer RNA molecules are packaged with higher selectivity than shorter ones. However, this trend did not hold across shuffle types. For example, the 472-nt SI insert exhibited a higher packaging fraction than the 3,570-nt SIV insert (**Fig. 4B**). These results indicate that RNA sequence, in addition to length, plays a significant role in determining selectivity.

The total amount of viral sequence within the insert does not appear to be a major determinant of selectivity. If it were, SI and SII would be expected to exhibit higher packaging fractions than SIII, SIV, and SV, as they contain larger fractions of viral sequence (and smaller fractions of shuffled sequence) (**Fig. 4A, right, “Shuffle (%)”**). However, the packaging fractions of SII inserts are consistently lower than those of SIII inserts across all lengths tested (**Fig. 4B**). These findings suggest that the total amount of viral sequence alone does not determine selectivity.

Likewise, the physical size of the insert transcripts does not appear to be a dominant factor in selectivity. SI and SII transcripts are predicted to adopt more compact structures (smaller MLDs) than SIII, SIV, and SV transcripts (**Fig. 4A, right, “MLD (bp)”**; and **SI Appendix, Fig. S11**). If compactness were a dominant factor, SI and SII would be expected to have higher packaging fractions than SIII, SIV, and SV. However, SII consistently shows lower packaging fractions than SIII across all lengths tested (**Fig. 4B**). These results suggest that physical size does not play a dominant role in selective packaging.

By contrast, the presence of packaging signal stem-loops has a clear and significant impact on selectivity. Consider two pairs of shuffle types: SI and SII, and SIII and SIV. Within each pair, the insert sequences are identical except that SI and SIII contain the proposed packaging signal stem-loops, while SII and SIV do not (**Fig. 4A, right, “Stem-loops”**). We observed that SI packaging fractions were approximately twice as high as those of SII, and SIII fractions were similarly about twice as high as those of SIV (**Fig. 4B**). This consistent trend across all lengths tested demonstrates that inserts containing packaging signal stem-loops achieve significantly higher selectivity, supporting the importance of these signals in the packaging process.

Quantitatively, we found that selectivity correlates more strongly with the number of packaging signal stem-loops than with the other properties tested. To assess these relationships, we calculated Pearson correlation coefficients (r) and p-values (p) for the packaging fraction as a function of the shuffle percentage, the MLD, and the number of stem-loops, using the full-length inserts shown in **Fig. 4A**. The correlation was strongest for the number of stem-loops (r=0.96, p=0.01) compared to shuffle percentage (r=-0.56, p=0.3) and MLD (r=-0.55, p=0.3) (**Fig. 4C**). These results align with the qualitative trends described above, further emphasizing the important role of stem-loops in packaging selectivity.

### Collective properties of the packaging signal stem-loops

To investigate potential collective properties of the packaging signal stem-loops, we generated additional shuffle types with varying numbers of these loops. Rather than varying loops at random, we focused on three specific loops that have received particular attention for their roles in packaging. These include the TR-loop, widely regarded as the preeminent feature in MS2 packaging (39), and two flanking stem-loops adjacent to TR in the MS2 genome, termed “TR-1” and “TR+1” (16) (**Fig. 5A**). These flanking loops have been proposed to work alongside TR to initiate packaging (25). We systematically added and removed these loops—either TR alone or TR with its flanking loops—to create four additional shuffle types, as shown in **Fig. 5A**:

**Fig. 5.**
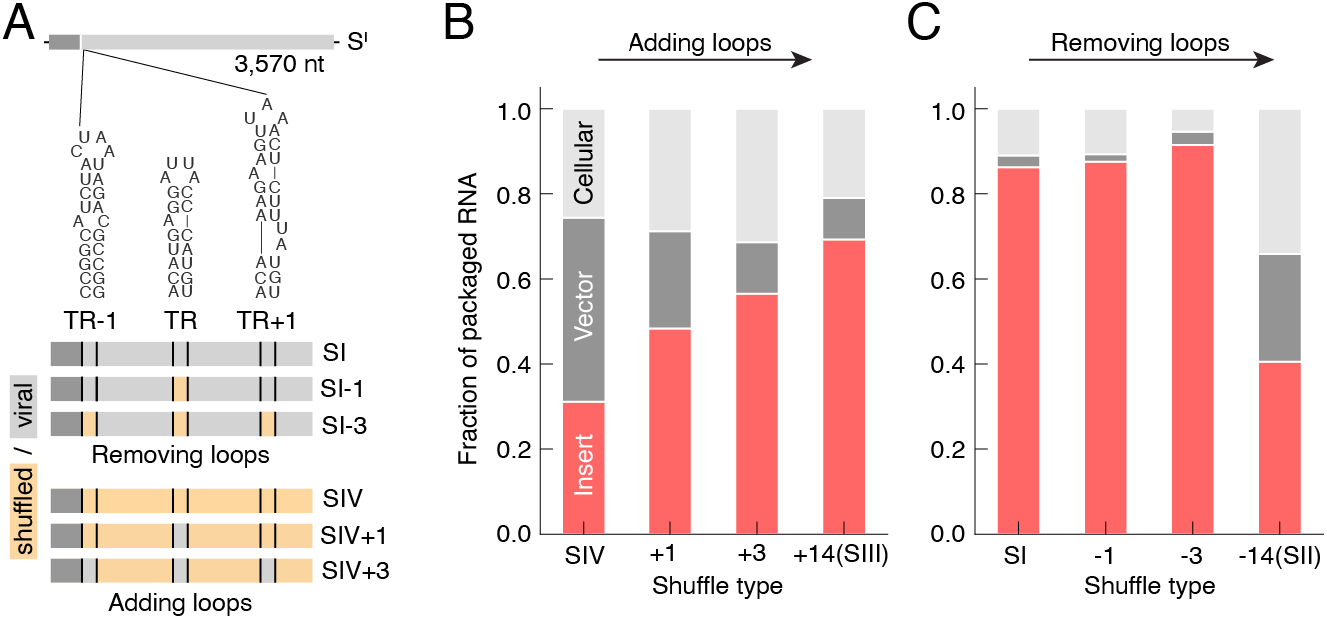
Selectivity depends on the number of stem-loops. **A**. Four new shuffle types (SI-1, SI-3, SIV+1, and SIV+3) were generated to investigate the roles of the TR loop and its flanking stem-loops (TR-1 and TR+1). For detailed descriptions of these shuffle types, see the **main text** and **SI Appendix, Dataset S1. B**. Adding TR and its flanking pair of loops (SIV+1 and SIV+3) to an otherwise random insert increases the packaging fraction of the insert transcripts dramatically, while adding an additional 11 loops (SIII) further increases the packaging fraction only modestly. **C**. Removing TR and the flanking loops (SI-1 and SI-3) does not significantly affect selectivity, whereas removing an additional 11 loops (SII) leads to a dramatic drop in selectivity.

- **SI-1**: the SI sequence with the TR-loop shuffled out.
- **SI-3**: SI with TR, TR-1, and TR+1 shuffled out.
- **SIV+1**: the SIV sequence with TR added in.
- **SIV+3**: SIV with TR, TR-1, and TR+1 added in.

Measurements of the packaging fractions for these new shuffle types reveal a non-trivial relationship between selectivity and the number of stem-loops. Consider the effect of adding stem-loops to the otherwise random SIV sequence: adding TR alone increased the packaging fraction from 31% to 48%, while adding the two flanking loops (TR-1 and TR+1) yielded an additional increase from 48% to 57% (**Fig. 5B**). Adding the remaining 11 loops required for a complete set (as in SIII) resulted in a further increase from 57% to 69% (**Fig. 5B**). These results show a clear positive correlation between packaging selectivity and the number of added stem-loops, but suggest diminishing returns with the addition of stem-loops beyond TR and its flanking loops.

Removing stem-loops further highlights this non-trivial relationship. When the TR loop is removed from the unshuffled SI insert, we observe no decrease in the packaging fraction, and further removal of the two flanking loops also yields no decrease (**Fig. 5B**). However, removing all 14 loops (as in SII) leads to a substantial decrease in the packaging fraction from 92% to 41% (**Fig. 5B**). These findings suggest that the absence of a single loop, such as the TR loop, or even a small subset of loops, does not significantly impair selectivity as long as other loops remain to compensate.

These results raise the question of whether additional stem-loops in the MS2 sequence might be important. A comparison of SI and SIII (**Fig. 4B**) indicates that the regions in between the 15 stem-loops in the MS2 sequence affect selectivity, but it is unclear if their effect is caused by additional stem-loops or some other properties. For example, it is possible that these regions provide a favorable context for the proper formation of the 15 stem-loop structures (40). Alternatively, the regions could provide a structural scaffold that positions the 15 stem-loops in special configurations that promote packaging.

We know that the configuration of the stem-loops can affect selectivity because we see differences in the packaging fractions of pMS2′ and SI (**Figs. 2A and 4B**). These inserts are circularly permuted homologs, meaning their transcripts contain the same set of stem-loop sequences but with different relative positions along the overall sequence. Notably, SI exhibits a lower packaging fraction (86 ± 3%, across duplicate biological experiments) compared to pMS2′ (97.4 ± 0.2%). This observation suggests that packaging signal stem-loops influence selectivity not only through their presence in the RNA sequence but also through their specific arrangement within the sequence.

Together, these results demonstrate that stem-loop-mediated packaging selectivity is a collective phenomenon driven by an ensemble of loops distributed throughout the RNA molecule. The contribution of any individual stem-loop to selectivity is difficult to quantify, as it depends on the presence and interplay of other loops within the molecule.

### Comparison to natural MS2

Finally, we compared the packaging fractions measured above to that of natural MS2. To determine the packaging fraction of natural MS2, we infected F+ *E. coli* cells with the virus, harvested the newly synthesized particles, extracted their packaged RNA, and sequenced it using the same protocols as before. We found that 99.6% of the sequencing reads from natural MS2 aligned to the MS2 genome, 0.2% aligned to the host genome, and 0.2% failed to align. The 99.6% packaging fraction of natural MS2 is similar in magnitude to the 97.4% value measured for the pMS2′ system. However, natural MS2 packages only 0.2% of cellular RNA, an order of magnitude less than the 2.2% packaged by pMS2′ particles (**Fig. 2A, right**). Thus, natural MS2 packages its own RNA with higher fidelity than our system does.

The near-perfect packaging fraction of natural MS2 is remarkable, given that MS2 coat proteins are highly promiscuous in vitro, packaging a broad range of viral and non-viral RNA molecules. Beyond foreign RNA, MS2 coat proteins can efficiently package synthetic polymers (41), metal nanoparticles (42), quantum dots (43), and even other proteins (44). Understanding how a process that is so permissive in vitro achieves such strict fidelity in vivo remains an open and intriguing question.

Unlike some eukaryotic RNA viruses, which sequester their subunits into membrane-enclosed “replication factories” (45) or membrane-less “viroplasms” (46), natural MS2 assembles directly in the cytoplasm amid a crowded mixture of host molecules (47). Achieving near-perfect packaging fidelity under these conditions likely requires multiple, complementary mechanisms involving multiple viral proteins. One possible mechanism is genome amplification by the viral replicase, which increases the abundance of viral RNA relative to host RNA, thereby enhancing its odds of being packaged (48). Another possible mechanism is genome labeling by the maturase protein, which binds to the 3′ end of the viral RNA and could mark it for encapsidation (49). Yet another possible mechanism is temporal regulation by the lysis protein (50), which ruptures the cell late in the replication cycle and could bias packaging toward RNA molecules that are more abundant early on (51).

These additional viral proteins—replicase, maturase, and lysis—are all absent from our simplified system, which expresses only the coat protein. While our system achieves high fidelity packaging using coat protein alone, achieving the near-perfect fidelity of natural MS2 could require the interplay of multiple viral components. Elucidating the collective effects of these components will be the subject of future work.

## Conclusions

By blocking expression of the viral replicase, maturase and lysis proteins, our experiments focus exclusively on whether—and to what extent—the coat proteins recognize specific RNA structures to achieve selective packaging. Through quantitative measurements of packaging fractions across a diverse set of RNA molecules—in which we systematically varied length, physical size, and the number of packaging signal stem-loops—we uncovered critical features of packaging selectivity that were inaccessible to previous approaches. We summarize our findings as follows:

- MS2 coat proteins alone are capable of selectively packaging MS2 RNA, achieving packaging fractions as high as 97% under the conditions tested (**Fig. 2**).
- RNA length influences packaging selectivity, but RNA sequence plays a more important role (**Fig. 3**).
- Specific groups of locally folded stem-loop structures distributed across the MS2 sequence have a disproportionately large effect on selectivity (**Fig. 4**).
- These packaging signal stem-loops function collectively, with no single loop—not even the famous TR loop—being strictly essential for selective packaging (**Fig. 5**).

These findings are qualitatively consistent with key aspects of the packaging signal hypothesis proposed by Stockley and coworkers (52), which posits that multiple copies of packaging signals in viral RNA molecules direct capsid assembly and facilitate RNA packaging. While this model has been highly influential in shaping current understanding of viral assembly (53), it has not yet been developed to the point of making quantitative predictions about packaging selectivity in cells. In some cases, the model fails to capture even qualitative trends observed in experiments (54). Our results provide a basis for refining this model and developing new ones that account for selectivity in quantitative detail (55). Bringing such models into quantitative agreement with experiment would not only deepen our understanding of plus-strand RNA viruses, but could also support the rational design of synthetic capsids that package defined RNA cargoes (56)—an important goal for emerging applications in gene delivery and genetic medicine (57).

## Materials and Methods

We prepared all buffer solutions using molecular biology-grade reagents and milli-Q water, and sterilized the solutions by autoclaving at 121°C for 20 minutes before use. The following buffers and media were used:

- **TE buffer**: 10 mM Tris-HCl (pH 7.0), 1 mM EDTA.
- **TNE buffer**: 50 mM Tris-HCl (pH 7.0), 100 mM NaCl, 1 mM EDTA.
- **LB media**: 10 g/L tryptone, 5 g/L yeast extract, 10 g/L sodium chloride, supplemented with either 0.05 g/L kanamycin or 0.10 g/L ampicillin.

### Plasmids

Plasmid constructs were prepared by Twist Bioscience. Inserts were chemically synthesized and then cloned into pET vectors (see **SI Appendix, Fig. S1**). The sequence of each construct was verified by next-generation sequencing, as reported in **SI Appendix, Dataset S1**.

### Cells

For packaging experiments, we used chemically competent *E. coli* strain BL21(DE3) (New England Biolabs). To produce natural MS2, we used F-pili-producing *E. coli* strain HS(pFamp)R (ATCC 700891).

### Insert sequence design

We generated all insert sequences used in this study by permuting, shuffling, and/or truncating portions of the pMS2′ sequence (**SI Appendix, Fig. S1**). To generate SI, we applied a circular permutation of pMS2′, placing the 5’-end of the insert at the start of the coat gene. The shuffling procedure used for SII–SIV preserves the overall dinucleotide frequencies across the insert (**SI Appendix, Fig. S9**). The Python code for these dinucleotide-preserving shuffles was developed by the Clote laboratory at Boston College (30). To generate the shuffled coat gene in SV, we used a shuffling procedure that preserves both codon and dinucleotide frequencies (**SI Appendix, Fig. S5**). The Python code for this shuffle was developed in our lab. We produced inserts of varying lengths by truncating the sequence at the 394th, 473rd, 1,329th, or 2,236th nucleotide (**Figs. 3 and 4**). Accordingly, the pcoat′ insert corresponds to the 394-nt truncation of SV.

### Structural analysis of the insert transcripts

We analyzed the structures of the insert transcripts using RNA folding models, including RNAfold and RNAsubopt from the ViennaRNA package (36). For each insert, we used RNAfold to predict the minimum free-energy secondary structure (see **SI Appendix, Fig. S10**) and RNAsubopt to predict an equilibrium ensemble of suboptimal secondary structures, sampled according to their Boltzmann weights. From these structures, we computed the maximum ladder distance of each transcript. The MLD was calculated as the mean across 1,000 suboptimal structures (see **Fig. 4A, right, “MLD (bp)”** and **SI Appendix, Fig. S11**). We used Python code developed by the Das laboratory at Stanford University (58) to compute the MLD of each secondary structure.

### In-cellulo packaging experiment

In parallel, we transformed *E. coli* strain BL21(DE3) with each of the packaging plasmids and streaked the transformed cells onto LB-agar plates containing kanamycin. A single colony was transferred to 100 mL of LB media containing kanamycin, and the liquid culture was incubated at 37°C in a 250 mL Erlenmeyer flask with shaking at 250 rpm. No IPTG was added. After 24 h, we lysed the cells and purified the virus-like particles from the cell debris and unpackaged nucleic acids. Our purification protocols are detailed in **SI Appendix, Supporting text**. Briefly, for RNA sequencing measurements, we performed multiple rounds of nuclease digestion using high concentrations of RNase A and DNase I to digest any unpackaged nucleic acids, followed by sucrose density centrifugation to separate the nuclease-resistant virus-like particles from the digested material. For microscopy measurements, we used a combination of sucrose density centrifugation and size-exclusion chromatography. The purity and yield of the resulting particles was measured using UV spectrophotometry, native agarose gel electrophoresis, and denaturing SDS-PAGE. The particles were stored in TNE buffer at -80°C prior to use.

### iSCAT microscopy

We determined the mass distribution of the purified virus-like particles using iSCAT microscopy. A 20-μL drop containing 1 nM particles diluted in water was added to a functionalized glass coverslip on a TwoMP iSCAT microscope (Refeyn). We functionalized the coverslips by treating them with poly-L-lysine, which imparts a positive charge to the surface, enhancing binding to the negatively charged particles. As the particles bind to the surface, the microscope records changes in intensity associated with each binding event, which are proportional to the mass of the particle. We recorded at least 2,000 particles per sample, with duplicate measurements performed for each.

To calibrate the iSCAT microscope we performed similar measurements on particles of known mass. Specifically, we used jackbean urease protein (90.8 kDa, Thermo Fisher), which forms clusters in solution, including 270-kDa trimers, 540-kDa hexamers, 820-kDa nonomers, and 1,080-kDa dodecamers. We also used self-assembled T=3 capsids of MS2 coat protein (90×13.7 kDa), which form 2,470-kDa particles, as well as natural MS2, which consists of 3,600-kDa particles of protein and RNA. Here, we note that, per unit mass, protein and RNA contribute equally to the iSCAT intensity (28, 59–61). The calibration measurements yielded a linear relationship between the measured intensity and the expected mass (see **SI Appendix, Fig. S3**). We used this linear relationship to infer the masses of the purified virus-like particles.

### Negative-stain electron microscopy

We prepared virus-like particles for TEM imaging by depositing a 6-μL drop of 10 nM particles in TNE buffer onto a clean piece of parafilm and placing a freshly glow-discharged formvar-carbon-coated 200-mesh copper grid (Electron Microscopy Sciences) onto the drop, carbon side down. After 2 min, we removed the grid from the drop and blotted it with filter paper. The grid was then placed onto a 10-μL drop of 1% uranyl acetate for 30 s, and blotted as before. Then, we placed the grid onto a fresh drop of 1% uranyl acetate for another 30 s, at which point the grid was blotted and left to dry completely. We imaged the grids using a Tecnai 12 electron microscope (FEI) with an accelerating voltage of 120 kV and a side-mount camera (AMT). Images were recorded at 97,000× magnification.

### Cryo-electron microscopy

We collected cryo-EM data using a Titan Krios G4i (Thermo-Fisher) operating at 300 keV with Bioquantum energy filter (slit width: 20 eV) and equipped with a K3 direct electron detector (Gatan). Micrographs were collected at 105,000× magnification (0.832 Å/pixel) by recording 50 frames over 1.93 seconds for a total dose of 50.18 e^-^/Å^2^. Data processing was carried out using Cryosparc v4.6.0 (62). Briefly, the dose-fractionated movies were subjected to motion correction and CTF estimation was carried out on resulting images. Particles were picked using blob picking followed by template picking. For pMS2′, a total of 728,471 particles were used for 3D refinement, and for pcoat′, a total of 313,055 particles were used. The overall map resolutions were estimated based on the gold-standard Fourier shell correlation (FSC 0.143) (63). The final maps were deposited into the Electron Microscopy Data Bank (EMD-48864, EMD-48865, EMD-48866, EMD-48867, and EMD-48868; see **SI Appendix, Table S2**). We generated initial models using ModelAngelo (64), using a combination of both sequence and non-sequence modes. Refinement was carried out using Phenix (65) and model adjustments were carried out in COOT (66). Models were deposited into the Protein Data Bank (PDB ID: 9N40 and 9N41; see **SI Appendix, Table S3**)

### RNA extraction

We extracted packaged RNA from 5 μg of purified particles using an RNeasy Mini kit (QIAGEN), following the manufacturer’s protocol. Total RNA was extracted from *E. coli* cells using the same kit, with the addition of RNAprotect bacteria reagent, according to the manufacturer’s protocol. We assessed the integrity of the extracted RNA using agarose gel electrophoresis, and the purity of the RNA with respect to protein contamination by UV spectrophotometry. RNA samples were stored in TE buffer at -80°C prior to sequencing.

### High-throughput RNA sequencing

We submitted 500 ng of extracted RNA in 30 μL of TE buffer for high-throughput RNA sequencing at the University of California, San Diego, Institute for Genomic Medicine Genomics Center. The RNA samples entered the standard sequencing pipeline just before the fragmentation step, ensuring that the sequencing libraries were generated without depletion of ribosomal RNA. Samples were sequenced on a NovaSeq 6000 S4 platform (Illumina), collecting at least 1 million 150-base-long paired-end reads per sample. The resulting reads, along with their associated quality metrics, were stored in FASTQ files for downstream analysis.

### Analysis of RNA sequencing data

Our RNAseq analysis protocol is detailed in **SI Appendix, Supporting text**. Briefly, we filtered the RNA sequencing reads for overall quality using fastp (v0.23.2) (67) with the following parameters: minimum read length of 15 (-l 15), sliding window size of 8 (-w 8), quality threshold of 15 (-q 15), maximum unqualified base percentage of 40% (-u 40), maximum number of ambiguous bases of 5 (-n 5), and adapter trimming (-a CTGTCTCTTATACACATCT). In our alignment protocol, the reads were treated as unpaired, meaning the paired-end nature of the data was not exploited. We used Bowtie2 (v2.4.5) (68) in local alignment mode (--local) to align up to one million reads per sample (-u 1000000) to the plasmid reference sequence, using default scoring settings for mismatches and gaps. Unaligned reads were captured (--un-gz) and subsequently mapped to the bacterial host genome using the same parameters. The alignment results, initially stored in SAM format, were converted to BAM format using samtools (v1.16.1) (69) view command. We then sorted the BAM files by genomic coordinates using samtools sort and computed coverage values using samtools mpileup. The resulting coverage values were used to calculate packaging fractions by integrating the coverage across the insert and then dividing by the total coverage. We report a summary of the aligned reads and packaging fractions in **SI Appendix, Table S1**.

### Amplification and purification of natural MS2

We amplified natural MS2 by infecting *E. coli* strain HS(pFamp)R with a stock sample of MS2 originally gifted to us by Peter Stockley at the University of Leeds, UK. We purified natural MS2 using the same protocols as those used for purifying virus-like particles, as detailed in **SI Appendix, Supporting text**.

## Acknowledgments

We thank Matt Mealka for helping us with size-exclusion chromatography, Ingrid Niesman for helping us with negative-stain TEM, Tommy Yiyang Zhou for helping us with RNAseq analysis, Fernando Vasquez for carrying out preliminary experiments, and Peter Stockley for supplying our initial stock of MS2. Some of the research was supported by the National Institute of General Medical Sciences of the National Institutes of Health (R35GM140803 to K.N.P. and R00GM127751 to R.F.G.) and the National Science Foundation (CAREER award number 2443955 to R.F.G.). RNA sequencing was conducted at the IGM Genomics Center, University of California, San Diego, using an Illumina NovaSeq 6000 that was purchased with funding from a National Institutes of Health SIG grant (S10OD026929). We thank the MSU RTSF Cryo-EM Core Facility for use of the Talos Arctica, and the California Metabolic Research Foundation for its support of biochemical research at San Diego State University. R.F.G acknowledges support from Ionis Pharmaceuticals.

## References

1. D. L. D. Caspar, A. Klug, Physical Principles in the Construction of Regular Viruses. Cold Spring Harb. Symp. Quant. Biol. 27, 1–24 (1962).

2. M. Comas-Garcia, Packaging of Genomic RNA in Positive-Sense Single-Stranded RNA Viruses: A Complex Story. Viruses 11, 253 (2019).

3. R. Milo, R. Phillips, Cell Biology by the Numbers (Garland Science, 2015).

4. A. Routh, T. Domitrovic, J. E. Johnson, Host RNAs, including transposons, are encapsidated by a eukaryotic single-stranded RNA virus. Proc. Natl. Acad. Sci. 109, 1907–1912 (2012).

5. K. Ghoshal, J. Theilmann, R. Reade, A. Maghodia, D. Rochon, Encapsidation of Host RNAs by Cucumber Necrosis Virus Coat Protein during both Agroinfiltration and Infection. J. Virol. 89, 10748–10761 (2015).

6. N. Shrestha, et al., Next generation sequencing reveals packaging of host RNAs by brome mosaic virus. Virus Res. 252, 82–90 (2018).

7. F. Qu, T. J. Morris, Encapsidation of turnip crinkle virus is defined by a specific packaging signal and RNA size. J. Virol. 71, 1428–1435 (1997).

8. T. Sugiyama, R. R. Hebert, K. A. Hartman, Ribonucleoprotein complexes formed between bacteriophage MS2 RNA and MS2 Protein in vitro. J. Mol. Biol. 25, 455–463 (1967).

9. H. E. Johansson, et al., A thermodynamic analysis of the sequence-specific binding of RNA by bacteriophage MS2 coat protein. Proc. Natl. Acad. Sci. 95, 9244–9249 (1998).

10. E. C. Hartman, et al., Quantitative characterization of all single amino acid variants of a viral capsid-based drug delivery vehicle. Nat. Commun. 9, 1385 (2018).

11. G. W. Witherell, J. M. Gott, O. C. Uhlenbeck, “Specific Interaction between RNA Phage Coat Proteins and RNA” in Progress in Nucleic Acid Research and Molecular Biology, W. E. Cohn, K. Moldave, Eds. (Academic Press, 1991), pp. 185–220.

12. G. G. Pickett, D. S. Peabody, Encapsidation of heterologous RNAs by bacteriophage MS2 coat protein. Nucleic Acids Res. 21, 4621–4626 (1993).

13. J. Carey, V. Cameron, P. L. De Haseth, O. C. Uhlenbeck, Sequence-specific interaction of R17 coat protein with its ribonucleic acid binding site. Biochemistry 22, 2601–2610 (1983).

14. R. I. Koning, et al., Asymmetric cryo-EM reconstruction of phage MS2 reveals genome structure in situ. Nat. Commun. 7, 12524 (2016).

15. X. Dai, et al., In situ structures of the genome and genome-delivery apparatus in a single-stranded RNA virus. Nature 541, 112–116 (2017).

16. Ó. Rolfsson, et al., Direct Evidence for Packaging Signal-Mediated Assembly of Bacteriophage MS2. J. Mol. Biol. 428, 431–448 (2016).

17. P. Simmonds, A. Tuplin, D. J. Evans, Detection of genome-scale ordered RNA structure (GORS) in genomes of positive-stranded RNA viruses: Implications for virus evolution and host persistence. RNA 10, 1337–1351 (2004).

18. D. Beckett, H.-N. Wu, O. C. Uhlenbeck, Roles of operator and non-operator RNA sequences in bacteriophage R17 capsid assembly. J. Mol. Biol. 204, 939–947 (1988).

19. A. M. Yoffe, et al., Predicting the sizes of large RNA molecules. Proc. Natl. Acad. Sci. 105, 16153–16158 (2008).

20. I. Kotta-Loizou, H. Peyret, K. Saunders, R. H. A. Coutts, G. P. Lomonossoff, Investigating the Biological Relevance of In Vitro-Identified Putative Packaging Signals at the 5′ Terminus of Satellite Tobacco Necrosis Virus 1 Genomic RNA. J. Virol. 93, 10.1128/jvi.02106-18 (2019).

21. K. Valegård, J. B. Murray, P. G. Stockley, N. J. Stonehouse, L. Liljas, Crystal structure of an RNA bacteriophage coat protein-operator complex. Nature 371, 623–626 (1994).

22. N. Patel, et al., Rewriting nature’s assembly manual for a ssRNA virus. Proc. Natl. Acad. Sci. 114, 12255–12260 (2017).

23. S. Shakeel, et al., Genomic RNA folding mediates assembly of human parechovirus. Nat. Commun. 8, 5 (2017).

24. R. S. Brown, D. G. Anastasakis, M. Hafner, M. Kielian, Multiple capsid protein binding sites mediate selective packaging of the alphavirus genomic RNA. Nat. Commun. 11, 4693 (2020).

25. R. Chandler-Bostock, et al., Genome-regulated Assembly of a ssRNA Virus May Also Prepare It for Infection. J. Mol. Biol. 434, 167797 (2022).

26. S. Tetter, et al., Evolution of a virus-like architecture and packaging mechanism in a repurposed bacterial protein. Science 372, 1220–1224 (2021).

27. S. Panahandeh, S. Li, B. Dragnea, R. Zandi, Virus Assembly Pathways Inside a Host Cell. ACS Nano 16, 317–327 (2022).

28. G. Young, et al., Quantitative mass imaging of single biological macromolecules. Science 360, 423–427 (2018).

29. L. Katz, C. B. Burge, Widespread Selection for Local RNA Secondary Structure in Coding Regions of Bacterial Genes. Genome Res. 13, 2042–2051 (2003).

30. P. Clote, Structural RNA has lower folding energy than random RNA of the same dinucleotide frequency. RNA 11, 578–591 (2005).

31. N. J. Atkinson, J. Witteveldt, D. J. Evans, P. Simmonds, The influence of CpG and UpA dinucleotide frequencies on RNA virus replication and characterization of the innate cellular pathways underlying virus attenuation and enhanced replication. Nucleic Acids Res. 42, 4527–4545 (2014).

32. A. P. Biela, A. Naskalska, F. Fatehi, R. Twarock, J. G. Heddle, Programmable polymorphism of a virus-like particle. Commun. Mater. 3, 1–9 (2022).

33. R. Twarock, R. J. Bingham, E. C. Dykeman, P. G. Stockley, A modelling paradigm for RNA virus assembly. Curr. Opin. Virol. 31, 74–81 (2018).

34. E. C. Dykeman, P. G. Stockley, R. Twarock, Solving a Levinthal’s paradox for virus assembly identifies a unique antiviral strategy. Proc. Natl. Acad. Sci. 111, 5361–5366 (2014).

35. A. Ben-Shaul, W. M. Gelbart, Viral ssRNAs Are Indeed Compact. Biophys. J. 108, 14–16 (2015).

36. R. Lorenz, et al., ViennaRNA Package 2.0. Algorithms Mol. Biol. 6, 26 (2011).

37. A. Borodavka, et al., Sizes of Long RNA Molecules Are Determined by the Branching Patterns of Their Secondary Structures. Biophys. J. 111, 2077–2085 (2016).

38. A. Gopal, et al., Viral RNAs Are Unusually Compact. PLOS ONE 9, e105875 (2014).

39. P. G. Stockley, et al., A simple, RNA-mediated allosteric switch controls the pathway to formation of a T= 3 viral capsid. J. Mol. Biol. 369, 541–552 (2007).

40. V. Bukina, A. Božič, Context-dependent structure formation of hairpin motifs in bacteriophage MS2 genomic RNA. Biophys. J. 123, 3397–3407 (2024).

41. T. Hohn, Role of RNA in the assembly process of bacteriophage fr. J. Mol. Biol. 43, 191–200 (1969).

42. S. L. Capehart, M. P. Coyle, J. E. Glasgow, M. B. Francis, Controlled Integration of Gold Nanoparticles and Organic Fluorophores Using Synthetically Modified MS2 Viral Capsids. J. Am. Chem. Soc. 135, 3011–3016 (2013).

43. C. E. Ashley, et al., Cell-Specific Delivery of Diverse Cargos by Bacteriophage MS2 Virus-like Particles. ACS Nano 5, 5729–5745 (2011).

44. J. E. Glasgow, S. L. Capehart, M. B. Francis, D. Tullman-Ercek, Osmolyte-Mediated Encapsulation of Proteins inside MS2 Viral Capsids. ACS Nano 6, 8658–8664 (2012).

45. J. A. den Boon, P. Ahlquist, Organelle-Like Membrane Compartmentalization of Positive-Strand RNA Virus Replication Factories. Annu. Rev. Microbiol. 64, 241–256 (2010).

46. G. Papa, A. Borodavka, U. Desselberger, Viroplasms: Assembly and Functions of Rotavirus Replication Factories. Viruses 13, 1349 (2021).

47. P. Pumpens, Single-Stranded RNA Phages: From Molecular Biology to Nanotechnology (CRC Press, 2020).

48. M. Eigen, C. K. Biebricher, M. Gebinoga, W. C. Gardiner, The hypercycle. Coupling of RNA and protein biosynthesis in the infection cycle of an RNA bacteriophage. Biochemistry 30, 11005–11018 (1991).

49. R. I. Koning, et al., Asymmetric cryo-EM reconstruction of phage MS2 reveals genome structure in situ. Nat. Commun. 7, 12524 (2016).

50. K. R. Chamakura, G. B. Edwards, R. Young, Mutational analysis of the MS2 lysis protein L. Microbiology 163, 961–969 (2017).

51. I. Mizrahi, R. Bruinsma, J. Rudnick, Packaging contests between viral RNA molecules and kinetic selectivity. PLOS Comput. Biol. 18, e1009913 (2022).

52. P. G. Stockley, et al., Packaging signals in single-stranded RNA viruses: nature’s alternative to a purely electrostatic assembly mechanism. J. Biol. Phys. 39, 277–287 (2013).

53. P. E. Prevelige, Follow the Yellow Brick Road: A Paradigm Shift in Virus Assembly. J. Mol. Biol. 428, 416–418 (2016).

54. Y. Song, et al., Limits of variation, specific infectivity, and genome packaging of massively recoded poliovirus genomes. Proc. Natl. Acad. Sci. 114, E8731–E8740 (2017).

55. J. D. Perlmutter, M. F. Hagan, The Role of Packaging Sites in Efficient and Specific Virus Assembly. J. Mol. Biol. 427, 2451–2467 (2015).

56. G. L. Butterfield, et al., Evolution of a designed protein assembly encapsulating its own RNA genome. Nature 552, 415–420 (2017).

57. A. Naskalska, J. G. Heddle, Virus-like particles derived from bacteriophage MS2 as antigen scaffolds and RNA protective shells. Nanomed. 19, 1103–1115.

58. H. K. Wayment-Steele, et al., Theoretical basis for stabilizing messenger RNA through secondary structure design. Nucleic Acids Res. 49, 10604–10617 (2021).

59. A. M. Goldfain, R. F. Garmann, Y. Jin, Y. Lahini, V. N. Manoharan, Dynamic Measurements of the Position, Orientation, and DNA Content of Individual Unlabeled Bacteriophages. J. Phys. Chem. B 120, 6130–6138 (2016).

60. R. F. Garmann, A. M. Goldfain, V. N. Manoharan, Measurements of the self-assembly kinetics of individual viral capsids around their RNA genome. Proc. Natl. Acad. Sci. 116, 22485–22490 (2019).

61. R. F. Garmann, et al., Single-particle studies of the effects of RNA–protein interactions on the self-assembly of RNA virus particles. Proc. Natl. Acad. Sci. 119, e2206292119 (2022).

62. A. Punjani, J. L. Rubinstein, D. J. Fleet, M. A. Brubaker, cryoSPARC: algorithms for rapid unsupervised cryo-EM structure determination. Nat. Methods 14, 290–296 (2017).

63. R. Henderson, et al., Outcome of the First Electron Microscopy Validation Task Force Meeting. Structure 20, 205–214 (2012).

64. K. Jamali, et al., Automated model building and protein identification in cryo-EM maps. Nature 628, 450–457 (2024).

65. D. Liebschner, et al., Macromolecular structure determination using X-rays, neutrons and electrons: recent developments in Phenix. Acta Crystallogr. Sect. Struct. Biol. 75, 861–877 (2019).

66. P. Emsley, K. Cowtan, Coot: model-building tools for molecular graphics. Acta Crystallogr. D Biol. Crystallogr. 60, 2126–2132 (2004).

67. S. Chen, Y. Zhou, Y. Chen, J. Gu, fastp: an ultra-fast all-in-one FASTQ preprocessor. Bioinforma. Oxf. Engl. 34, i884–i890 (2018).

68. B. Langmead, S. L. Salzberg, Fast gapped-read alignment with Bowtie 2. Nat. Methods 9, 357–359 (2012).

69. P. Danecek, et al., Twelve years of SAMtools and BCFtools. GigaScience 10, giab008 (2021).

70. R. F. Garmann, M. Comas-Garcia, A. Gopal, C. M. Knobler, W. M. Gelbart, The assembly pathway of an icosahedral single-stranded RNA virus depends on the strength of inter-subunit attractions. J. Mol. Biol. 426, 1050–1060 (2014).

