## Supporting Information for "Measuring the selective packaging of RNA molecules by viral coat proteins in cells"

**This PDF file includes:**

Supporting text
Figures S1 to S11
Table S1
Legends for Movies S1
Legends for Datasets S1 to S2
SI References

### Supporting Information Text

#### Methods

**Details of the purification of virus-like particles.** After 24 hours of incubation, we collected the cells by spinning the liquid culture at 10,000 rpm for 30 minutes at 4°C in a Fiberlite F21-8x50y rotor (Thermo Scientific) and discarding the supernatant. The pellet was then resuspended in 25 mL of TNE buffer and mechanically lysed by sonication. The system was subjected to six intervals of 20 seconds at 20 kHz using a 120W sonicator (Fisher Scientific) operated at 20% amplitude with a CL-18 probe (Qsonica). Large cellular debris was removed by spinning the lysate at 12,000 rpm for 50 minutes at 4°C in a Fiberlite F21-8x50y rotor. Lipids were removed by chloroform extraction. We matched the sample volume with an equal volume of chloroform, shook vigorously for 3 minutes, and then spun the emulsion at 3,600 rpm for 10 minutes at 4°C in a SX4750 swinging bucket rotor (Beckman Coulter). The top aqueous layer was removed and added to fresh chloroform, and the process was repeated until the white interface between layers became vanishingly thin. From here on, the sample was subjected to different purification protocols depending on whether it was going to be analyzed by RNA sequencing experiments or microscopy experiments:

**Sequencing samples.** A sample destined for RNA sequencing experiments was subjected to two rounds of sucrose density centrifugation, with nuclease treatments before and after each round. The goal was to completely remove any unpackaged RNA or DNA that was not protected by a coat-protein capsid, ensuring that the final sequencing results reflected only the composition of the packaged RNA.

Directly after the chloroform extraction step, we passed the sample through a 0.2-μm PES syringe filter and froze it at -80°C. After thawing at room temperature, the sample was treated with 5 μg/mL of RNase A overnight at room temperature. Next, we performed a sucrose cushion separation by layering 12 mL of sample on top of 1 mL of 15% (wt/vol) sucrose in TNE buffer, and spun the system at 36,000 rpm for 6 hours at 4°C in a SW 41 rotor (Beckman Coulter). After spinning, the supernatant was decanted, and the pellets were resuspended in 0.5 mL of TNE buffer containing 5 μg/mL RNase A (Invitrogen) and 40 U/mL DNase I (Thermo Scientific). The sample was incubated at 37°C for 1 hour, and then left at room temperature overnight. The next day, we performed a second sucrose cushion separation by layering 10 mL of sample over 2 mL of 20% sucrose in TNE buffer, followed by another spin, as before. The resulting pellet was resuspended in 0.5 mL of TNE buffer containing 5 μg/mL RNase A and 40 U/mL DNase I and incubated at 37°C for 1 hour. The sample was then subjected to 5 rounds of centrifugal filtration, in which 0.5 mL of sample was added to a 0.5-mL 100-kDa centrifugal filter unit (MilliporeSigma), spun at 10,000 rcf for 5 minutes at room temperature, and resuspended in 0.5 mL of fresh TNE buffer. After 5 rounds, the sample was collected, and its purity and concentration were determined by gel electrophoresis and UV spectrophotometry, assuming an extinction coefficient of 8.03 mL $mg^{-1}$ $cm^{-1}$ at 260 nm (60), as shown in **Fig. S2**.

**Microscopy samples.** A sample destined for iSCAT and electron microscopy experiments was subjected to purification by sucrose cushion, sucrose gradient, and size-exclusion chromatography, without nuclease treatment. While MS2 virus-like particles are resistant to nuclease digestion, prolonged treatments—such as those used to prepare samples for RNA sequencing experiments—could affect the packaged RNA cargo. To preserve the integrity of the RNA for structural characterization, we omitted the nuclease treatment steps in favor of additional separation steps.

Immediately following chloroform extraction, we performed a sucrose cushion separation as described earlier by layering 12 mL of sample on top of 1 mL of 15% (wt/vol) sucrose in TNE buffer and spinning at 36,000 rpm for 6 hours at 4°C in a SW 41 rotor. After spinning, the supernatant was decanted, and the pellet was resuspended in 0.5 mL of TNE buffer. We then performed a sucrose gradient separation by layering 1 mL of the resuspended material on top of a 12 mL 10-40% sucrose gradient prepared in TNE buffer according to previously published protocols (70). The sample was spun at 36,000 rpm for 2 hours in a SW 41 rotor, and the central band was collected by puncturing the thin-wall ultraclear centrifuge tube (Beckman) from the side and aspirating the sample using a needle and syringe. The collected sample was dialyzed against fresh TNE buffer and then fractionated by size-exclusion chromatography using an AKTA go FPLC system with a HiLoad 16/600 Superdex 200 pg column (Cytiva) that was pre-equilibrated with TNE buffer and operated at a flow rate of 1.0 mL/min and a pressure of 1.5 MPa. The collected fractions were analyzed by SDS-PAGE, and the fractions containing pure MS2 coat protein were pooled together. Finally, we concentrated the pooled sample to approximately 5 mg/mL using a 100-kDa centrifugal filter unit (MilliporeSigma).

**Details of the production of natural MS2.** We prepared natural MS2 by infecting F+ *E. coli* cells with a stock sample of MS2 originally provided by Peter Stockley at the University of Leeds, UK. To initiate the culture, we added 0.1 mL of a frozen glycerol stock of *E. coli* strain HS(pFamp)R to 50 mL of LB broth containing ampicillin. The culture was incubated at 37°C in a 250 mL Erlenmeyer flask with shaking at 250 rpm until the optical density at 600 nm reached 0.4–0.5. At this point, we added $10^7$ plaque-forming units of MS2, yielding a multiplicity of infection of approximately $10^{-3}$. After 24 h of incubation with shaking at 37°C, we spun the culture at 10,000 rpm for 30 minutes in a Fiberlite F21-8x50y rotor to pellet uninfected cells and cell debris. The supernatant, which contained the MS2 particles, was collected and purified following the protocols described above.

**RNAseq data analysis.** Below, we outline the general RNA sequencing analysis pipeline used in our study and we provide high-level example code that illustrates the key steps:

1. **Reference index generation.** We combined the insert and vector sequences into a single fasta file. Then we used samtools faidx (v1.16.1) (69) to index the fasta file, and bowtie2-build (v2.4.5) (68) to create the index for alignment. This process was repeated for the host genome.

```
# Generate FASTA index using samtools
samtools faidx my_construct.fasta

# Build Bowtie2 index files from the FASTA
bowtie2-build my_construct.fasta my_construct
```

2. **Read filtering with fastp.** We used fastp (v0.23.2) (67) to filter low-quality bases, trim adapters, and discard short reads. Filtering parameters included a minimum read length of 15 (-l 15), sliding window size of 8 (-w 8), quality threshold of 15 (-q 15), maximum unqualified base percentage of 40% (-u 40), maximum of 5 ambiguous bases (-n 5), and specific adapter trimming (-a CTGTCTCTTATACACATCT). Only unpaired reads (R1) were analyzed.

```
# Filter unpaired reads for overall quality
fastp \
  -i raw_reads_R1.fastq \
  -o filtered_reads_R1.fastq \
  -l 15 \
  -w 8 \
  -q 15 \
  -u 40 \
  -n 5 \
  -a CTGTCTCTTATACACATCT
```

3. **Bowtie2 alignment.** We used Bowtie2 in local mode to align up to one million filtered reads per sample to the plasmid reference. Unaligned reads were subsequently aligned to the host reference. Alignments were performed with 8 threads using default scoring parameters.

```
# Align one million filtered reads to plasmid
bowtie2 --local -u 1000000 -p 8 -x plasmid_index \
  -U filtered_reads_R1.fastq \
  --un-gz unaligned_reads.gz \
  | samtools view -buS - \
  | samtools sort -o plasmid_aligned.bam

# Align any unaligned reads and align them to host
bowtie2 --local -u 1000000 -p 8 -x host_index \
  -U unaligned_reads.gz \
  | samtools view -buS - \
  | samtools sort -o host_aligned.bam
```

4. **SAM-to-BAM Conversion and Sorting.** We used samtools view to convert the alignment output from the SAM format to a compressed BAM format, and then we used samtools sort to sort by genomic coordinates.

```
# Convert and sort by genomic coordinate
samtools view -buS plasmid_aligned.sam \
  | samtools sort -o plasmid_aligned.bam
```

5. **Coverage Calculation and Packaging Fractions.** We used samtools mpileup to compute the per-base coverage, enabling coverage output for all positions (-aa) and allowing up to one million reads per position (-d 1000000).

```
samtools mpileup -aa -d 1000000 \
  -f my_construct.fasta \
  plasmid_aligned.bam \
  > plasmid_pileup.txt

samtools mpileup -aa -d 1000000 \
  -f host_reference.fasta \
  host_aligned.bam \
  > host_pileup.txt
```

We then integrated the coverage values across the insert region (and any vector or host regions) to determine the fraction of reads mapping to each. This analysis enabled us to calculate packaging fractions and assess the packaging selectivity for each experimental condition.

**Fraction of particles containing an insert transcript**

If we assume that each particle contains at most one copy of the insert transcript, then we can estimate the fraction of particles $F$ that contain the transcript as

$$F = (N/L) \cdot f,$$

where $N$ is the average number of nucleotides packaged per particle (measured by iSCAT), $L$ is the length of the insert transcript (measured in nucleotides), and $f$ is the packaging fraction (measured by RNAseq). Note this estimate represents an upper limit on the true fraction, since multiple inserts could be packaged into the same particle.

For the pcoat′ insert, which has $N = 3300\,nt$, $L = 394\,nt$, and $f = 0.03$, we calculate $F = 0.25$. Thus, we conclude that, at most, 1 out of every 4 particles contains a pcoat′ transcript.

A

1 GGGUGGGACCCCUUUCGGGGUCCUGCUCAACUUCCUGUCGAGCUAAUGCCAUUUUUAAUGUCUUUAGCGAGACGCUACCAUGGCUAUCGCUGUAGGUAGC 100
101 CGGAAUUCCAUUCCUAGGAGGUUUGACCUGUGCGAGCUU[UU/AA]AGUACCCUUGAUAGGGAGAACGAGACCUUCGUCCCCUCCGUUCGCGUUUACGCGGACGG 200
201 UGAGACUGAAGAUAACUCAUUCUCUUUAAAAUAUCGUUCGAACUGGACUCCCGGUCGUUUUAACUCGACUGGGGCCAAAACGAAACAGUGGCACUACCCC 300
301 UCUCCGUAUUCACGGGGGGCGUUAAGUGUCACAUCGAUAGAUCAAGGUGCCUACAAGCGAAGUGGGUCAUCGUGGGGUCGCCCGUACGAGGAGAAAGCCG 400
401 GUUUCGGCUUCUCCCUCGACGCACGCUCCUGCUACAGCCUCUUCCCUGUAAGCCAAAACUUGACUUACAUCGAAGUGCCGCAGAACGUUGCGAACCGGGC 500
501 GUCGACCGAAGUCCUGCAAAAGGUCACCCAGGGUAAUUUUAACCUUGGUGUUGCUUUAGCAGAGGCCAGGUCGACAGCCUCACAACUCGCGACGCAAACC 600
601 AUUGCGCUCGUGAAGGCGUACACUGCCGCUCGUCGCGGUAAUUGGCGCCAGGCGCUCCGCUACCUUGCCCUAAACGAAGAUCGAAAGUUUCGAUCAAAAC 700
701 ACGUGGCCGGCAGGUGGUUGGAGUUGCAGUUCGGUUGGUUACCACUAAUGAGUGAUAUCCAGGGUGCAUAUGAGAUGCUUACGAAGGUUCACCUUCAAGA 800
801 GUUUCUUCCUAUGAGAGCCGUACGUCAGGUCGGUACUAACAUCAAGUUAGAUGGCCGUCUGUCGUAUCCAGCUGCAAACUUCCAGACAACGUGCAACAUA 900
901 UCGCGACGUAUCGUGAUAUGGUUUUACAUAAACGAUGCACGUUUGGCAUGGUUGUCGUCUCUAGGUAUCUUGAACCCACUAGGUAUAGUGUGGGAAAAGG 1000
1001 UGCCUUUCUCAUUCGUUGUCGACUGGCUCCUACCUGUAGGUAACAUGCUCGAGGGCCUUACGGCCCCCGUGGGAUGCUCCUACAUGUCAGGAACAGUUAC 1100
1101 UGACGUAAUAACGGGUGAGUCCAUCAUAAGCGUUGACGCUCCCUACGGGUGGACUGUGGAGAGACAGGGCACUGCUAAGGCCCAAAUCUCAGCCAUGCAU 1200
1201 CGAGGGGUACAAUCCGUAUGGCCAACAACUGGCGCGUACGUAAAGUCUCCUUUCUCGAUGGUCCAUACCUUAGAUGCGUUAGCAUUAAUCAGGCAACGGC 1300
1301 UCUCUAGAUAGAGCCCUCAACCGGAGUUUGAAGAAUGGCUUCU[A/U]A[C/A]UUUACUCAGUUCGUUCUCGUCGACAAUGGCGGAACUGGCGACGUGACUGUCGCC 1400
1401 CCAAGCAACUUCGCUAACGGGGUCGCUGAAUGGAUCAGCUCUAACUCGCGUUCACAGGCUUACAAAGUAACCUGUAGCGUUCGUCAGAGCUCUGCGCAGA 1500
1501 AUCGCAAAUACACCAUCAAAGUCGAGGUGCCUAAAGUGGCAACCCAGACUGUUGGUGGUGUAGAGCUUCCUGUAGCCGCAUGGCGUUCGUACUUAAAUAU 1600
1601 GGAACUAACCAUUCCAAUUUUCGCUACGAAUUCCGACUGCGAGCUUAUUGUUAAGGCAAUGCAAGGUCUCCUAAAAGA[U/C]GGAAACCCGAUUCCCUCAGCA 1700
1701 AUCGCAGCAAACUCCGGCAUCUACUAAUAGACGCCGGCCAUUCAAACAUGAGGAUUACCCAUGUC[GA/UU]A[G/A]ACAACAAAGAAGUUCAACUCUUUAUGUAUUG 1800
1801 A[G/U]U[A/U]AACUCGCGAUCUUUCUCUCGAAAUUUACCAAUCAAUUGCUUCUGUCGCUACUGGAAGCGGUGAUCCGCACAGUGACGACUUUACAGCAAUUGCUUA 1900
1901 CUUAAGGGACGAAUUGCUCACAAAGCAUCCGACCUUAGGUUCUGGUAAUGACGAGGCGACCCGUCGUACCUUAGCUAUCGCUAAGCUACGGGAGGCGAAU 2000
2001 GGUGAUCGCGGUCAGAUAAAUAGAGAAGGUUUCUUACAUGACAAAUCCUUGUCAUGGGAUCCGGAUGUUUUACAAACCAGCAUCCGUAGCCUUAUUGGCA 2100
2101 ACCUCCUCUCUGGCUACCGAUCGUCGUUGUUUGGGCAAUGCACGUUCUCCAACGGUGCUCCUAUGGGGCACAAGUUGCAGGAUGCAGCGCCUUACAAGAA 2200
2201 GUUCGCUGAACAAGCAACCGUUACCCCCCGCGCUCUGAGAGCGGCUCUAUUGGUCCGAGACCAAUGUGCGCCGUGGAUCAGACACGCGGUCCGCUAUAAC 2300
2301 GAGUCAUAUGAAUUUAGGCUCGUUGUAGGGAACGGAGUGUUUACAGUUCCGAAGAAUAAUAAAAUAGAUCGGGCUGCCUGUAAGGAGCCUGAUAUGAAUA 2400
2401 UGUACCUCCAGAAAGGGGUCGGUGCUUUCAUCAGACGCCGGCUCAAAUCCGUUGGUAUAGACCUGAAUGAUCAAUCGAUCAACCAGCGUCUGGCUCAGCA 2500
2501 GGGCAGCGUAGAUGGUUCGCUUGCGACGAUAGACUUAUCGUCUGCAUCCGAUUCCAUCUCCGAUCGCCUGGUGUGGAGUUUUCUCCCACCAGAGCUAUAU 2600
2601 UCAUAUCUCGAUCGUAUCCGCUCACACUACGGAAUCGUAGAUGGCGAGACGAUACGAUGGGAACUAUUUUCCACAAUGGGAAAUGGGUUCACAUUUGAGC 2700
2701 UAGAGUCCAUGAUAUUCUGGGCAAUAGUCAAAGCGACCCAAAUCCAUUUUGGUAACGCCGGAACCAUAGGCAUCUACGGGGACGAUAUUAUAUGUCCCAG 2800
2801 UGAGAUUGCACCCCGUGUGCUAGAGGCACUUGCCUACUACGGUUUUAAACCGAAUCUUCGUAAAACGUUCGUGUCCGGGCUCUUUCGCGAGAGCUGCGGC 2900
2901 GCGCACUUUUACCGUGGUGUCGAUGUCAAACCGUUUUACAUCAAGAAACCUGUUGACAAUCUCUUCGCCCUGAUGCUGAUAUUAAAUCGGCUACGGGGUU 3000
3001 GGGGAGUUGUCGGAGGUAUGUCAGAUCCACGCCUCUAUAAGGUGUGGGUACGGCUCUCCUCCCAGGUGCCUUCGAUGUUCUUCGGUGGGACGGACCUCGC 3100
3101 UGCCGACUACUACGUAGUCAGCCCGCCUACGGCAGUCUCGGUAUACACCAAGACUCCGUACGGGCGGCUGCUCGCGGAUACCCGUACCUCGGGUUUCCGU 3200
3201 CUUGCUCGUAUCGCUCGAGAACGCAAGUUCUUCAGCGAAAAGCACGACAGUGGUCGCUACAUAGCGUGGUUCCAUACUGGAGGUGAAAUCACCGACAGCA 3300
3301 UGAAGUCCGCCGGCGUGCGCGUUAUACGCACUUCGGAGUGGCUAACGCCGGUUCCCACAUUCCCUCAGGAGUGUGGGCCAGCGAGCUCUCCUCGGUAGCU 3400
3401 GACCGAGGGACCCCCGUAAACGGGGUGGGUGUGCUCGAAAGAGCACGGGUGCGAAAGCGGUCCGGCUCCACCGAAAGGUGGGCGGGCUUCGGCCCAGGGA 3500
3501 CCUCCCCCUAAAGAGAGGACCCGGGAUUCUCCCGAUUUGGUAACUAGCUGCUUGGCUAGUUACCACCCAAGCUU

B

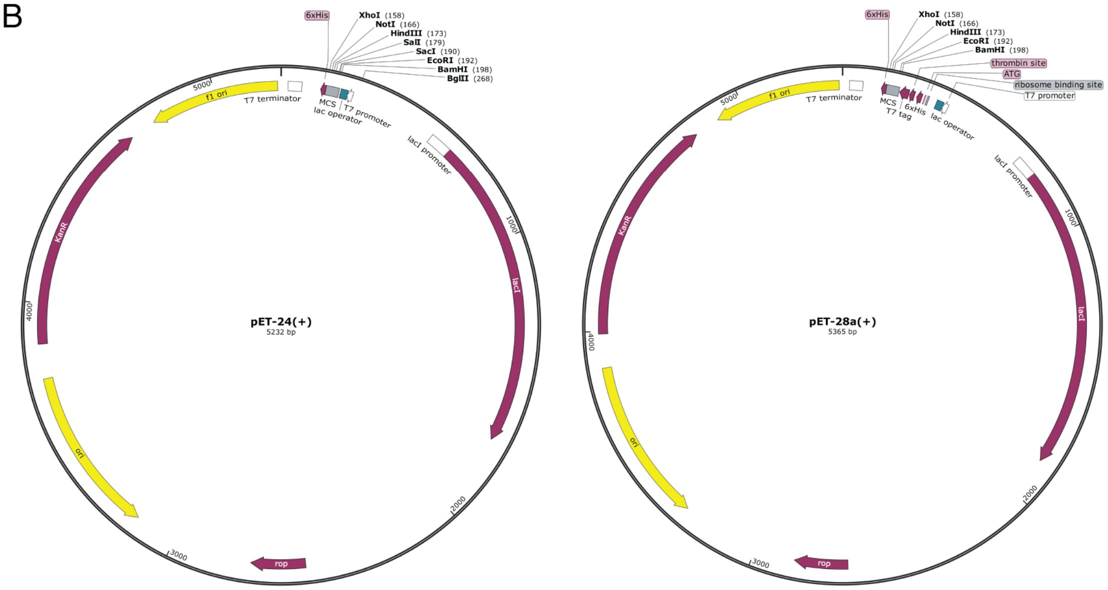


**Fig. S1. pMS2′ insert design. A.** We created the pMS2′ insert by introducing mutations into the wild-type MS2 genome sequence. Mutations are marked with solid boxes, with the wild-type sequence on top and the mutant sequence on bottom: the red box highlights an F4* mutation in the maturase gene (disabling maturase production); gray boxes denote N4* mutations in the coat gene (disabling coat production and packaging); the black box indicates an M1T mutation in the lysis gene (disabling lysis); and the blue boxes show a K3* mutation in the replicase gene (disabling replication) that also adds an additional stop codon in the lysis gene and maintains base pairing in the TR+1 stem-loop. We also included a HindIII restriction site at the 3' end for future in-vitro experiments. **B.** Diagram of the pET-24(+) vector used for pMS2′ and the pET-28a(+) vector used for other inserts. For the inserts placed in the pET-28a(+) vector, we removed the first 4 nts of the sequence to prevent overlap of the start codon. Full plasmid sequences, verified via next-generation sequencing, are provided in **Dataset S1**.

A

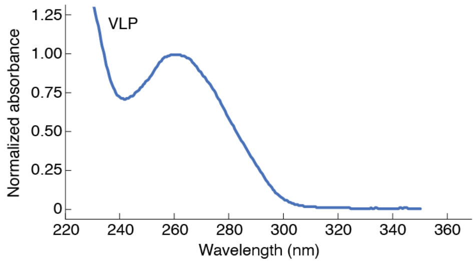


B

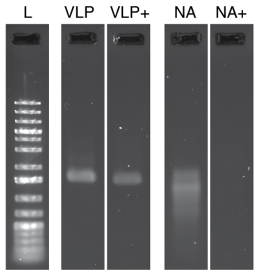


**Fig. S2. pMS2′ virus-like particles package RNA. A.** The UV absorbance spectrum of purified pMS2′ virus-like particles (VLP) shows a dominant peak at 260 nm, indicating the presence of nucleic acid. **B.** Agarose gel electrophoresis of pMS2′ particles (VLP) and nucleic acid extracted from these particles (NA), before and after treatment with RNase A (+). Extracted nucleic acid was isolated using phenol:chloroform. Ethidium-staining shows complete degradation of the extracted nucleic acid by RNase, indicating that it is RNA. Nucleic acid in the VLP is not digested by nuclease protected by the capsid.

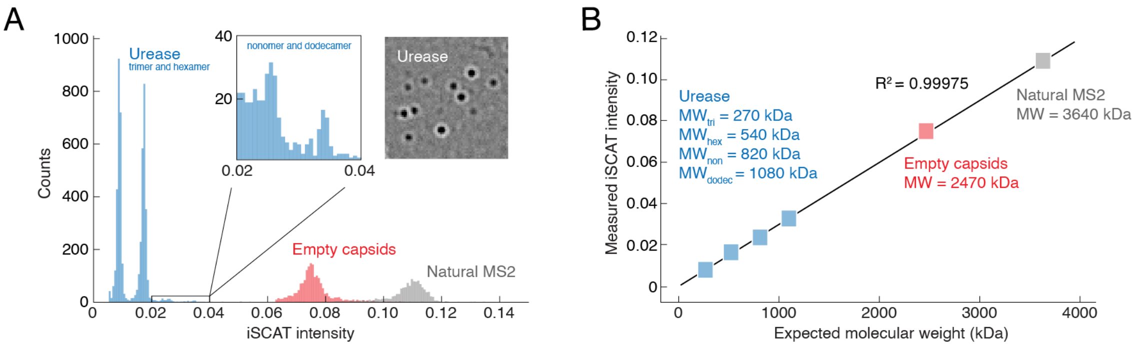


**Fig. S3. iSCAT experiments. A.** Histograms of iSCAT intensities for three mass standards: jackbean urease, empty MS2 capsids, and natural MS2 particles. Urease proteins form clusters of discrete sizes, while empty capsids and natural MS2 are monodisperse. The inset shows additional peaks in the urease histogram and a representative image of urease clusters binding to the coverslip. **B.** iSCAT intensity is linearly proportional to molecular weight, even for particles containing both protein and RNA, such as natural MS2.

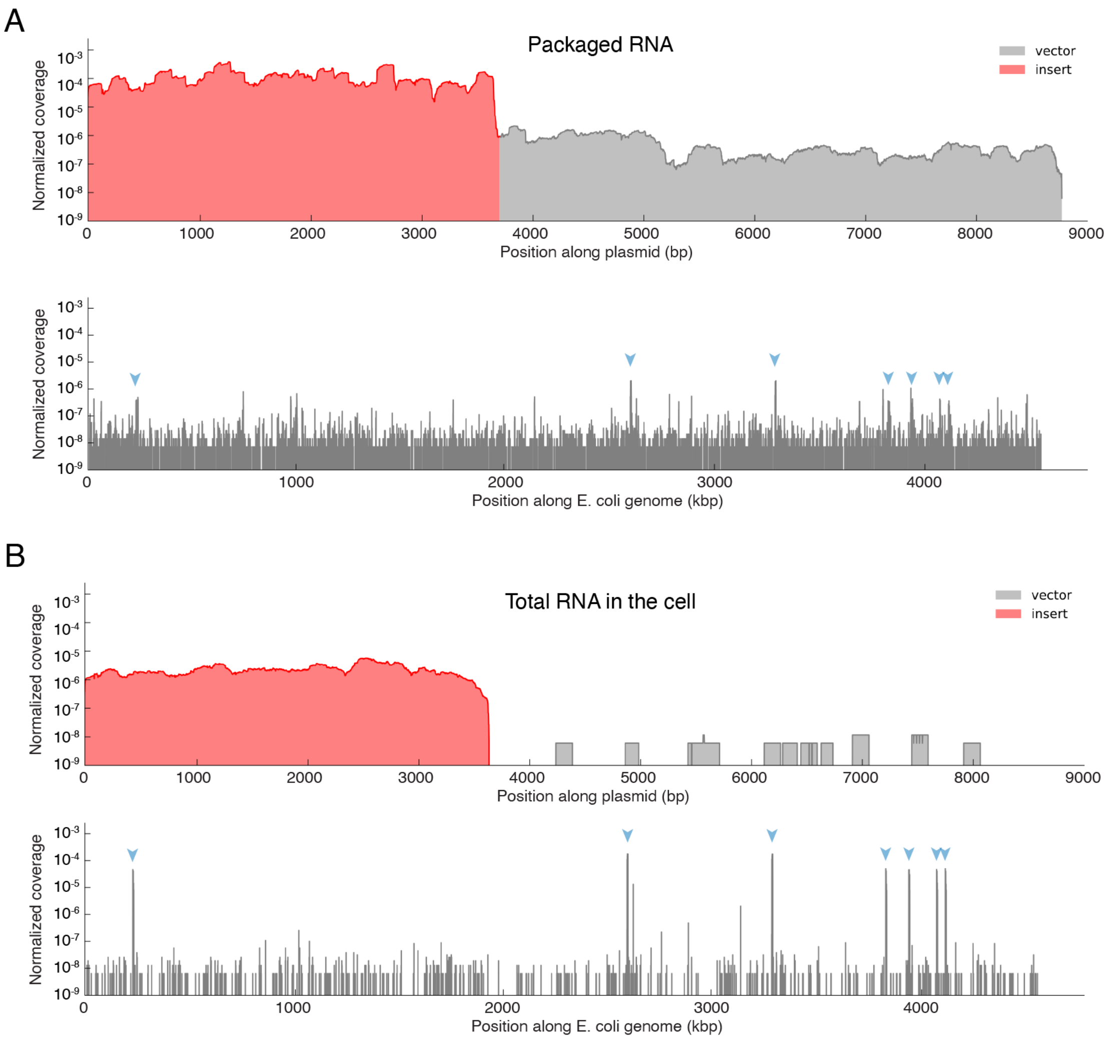


**Fig. S4. Sequencing packaged and total cellular RNA. A.** Coverage plot for RNA sequencing reads from packaged RNA in the pMS2′ packaging experiment. Insert coverage is shown in red, vector coverage in light gray, and host genome coverage in dark gray. Blue arrowheads mark ribosomal RNA genes. Most of the packaged RNA corresponds to the pMS2′ insert, while ribosomal transcripts are the predominant host RNA species. **B.** Coverage plot for total cellular RNA when packaging is abolished by a premature stop codon in the coat gene. Reduced insert coverage and increased ribosomal RNA coverage reflect the dominance of ribosomal transcripts in the cell. For each experiment, we recorded 100M reads.

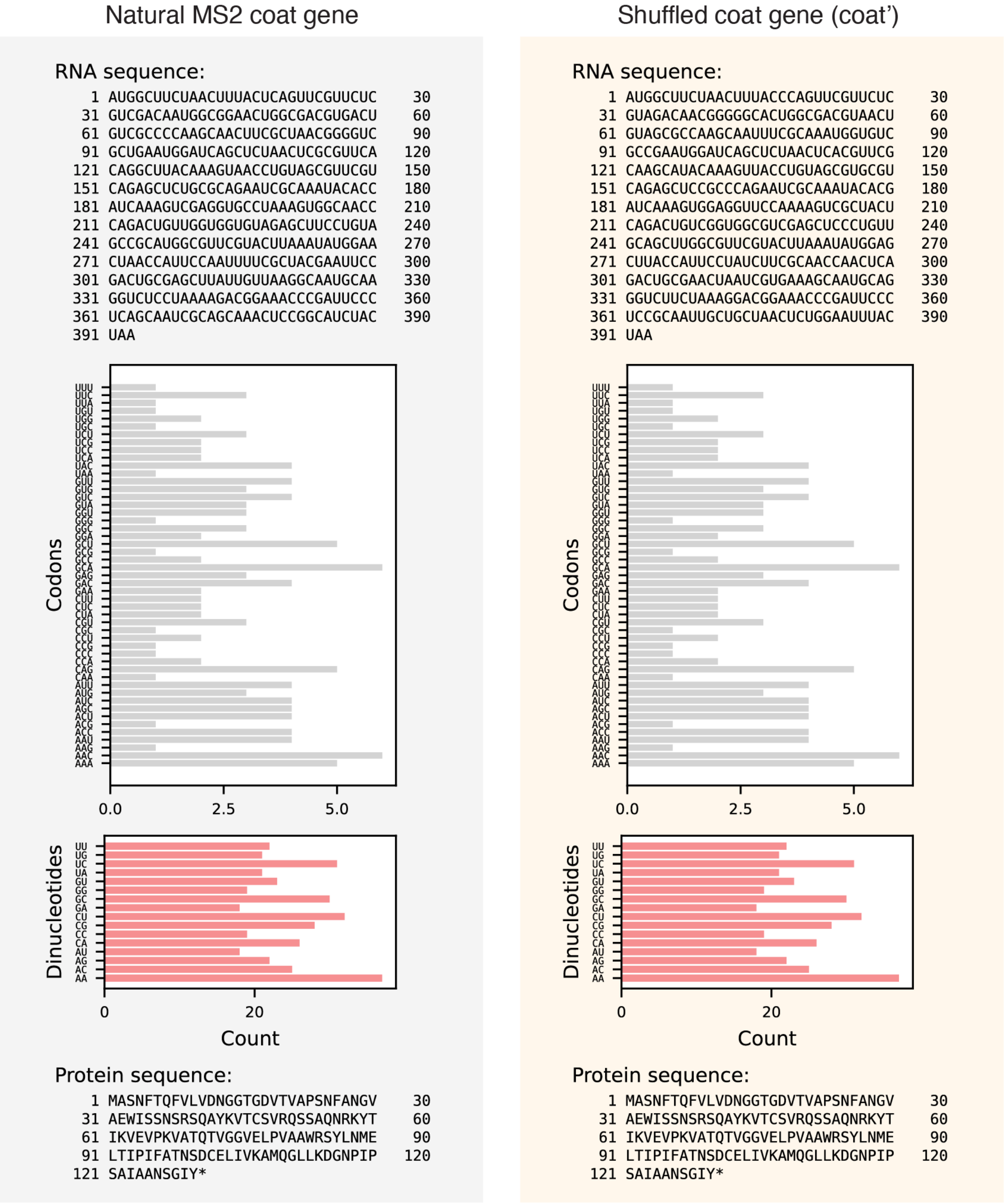


**Fig. S5. Comparison of natural and shuffled coat protein genes.** Comparison of the natural coat protein gene (used in pMS2′ and SI–SIV inserts) with the shuffled coat protein gene (used in pcoat′ and SV inserts). **B.** The shuffled gene differs at 58 positions but maintains identical codon and dinucleotide usage, as well as the same amino acid sequence.

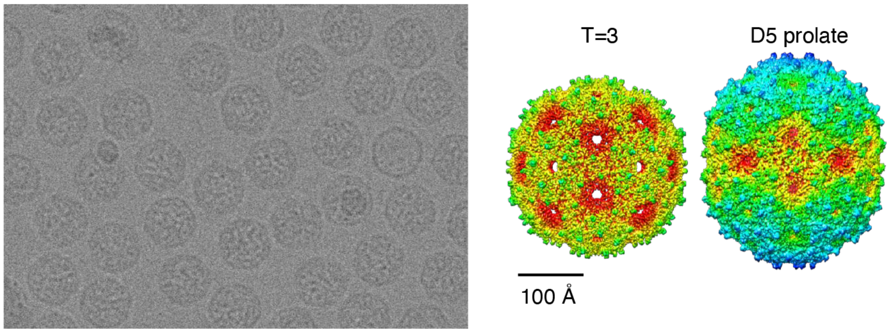


**Fig. S6. Cryo-EM of MS2' particles. Left.** Raw cryo-micrograph of the particles. **Right.** 3D reconstructions of the particles. Most of the particles have a T=3 structure, and some have a prolate shape with D5 symmetry (see **Tables S2 and S3** for details).

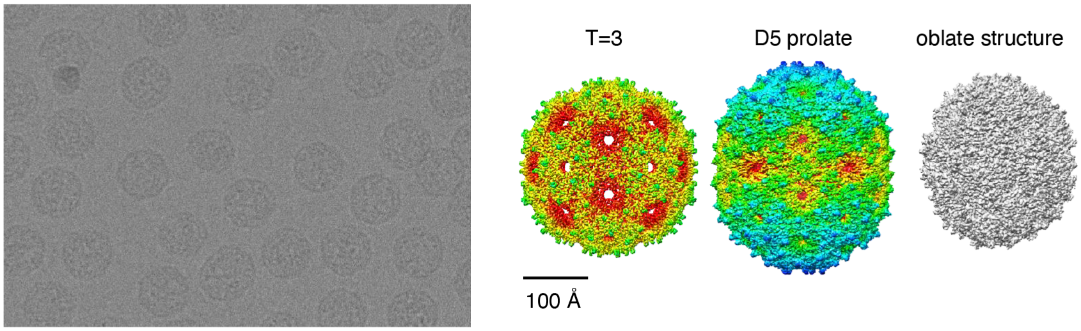


**Fig. S7. Cryo-EM of coat' particles. Left.** Raw cryo-micrograph of the particles. **Right.** 3D reconstructions of the particles. Most of the particles have a T=3 structure, some have a prolate shape with D5 symmetry, and others have an asymmetric oblate shape (see **Tables S2 and S3**, and **Movie S1** for details).

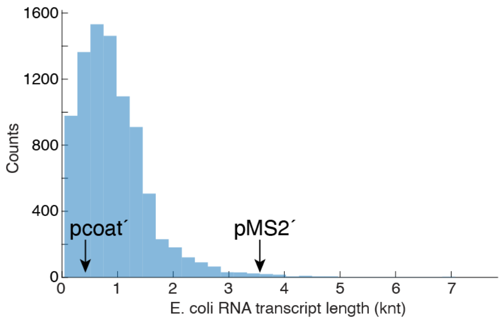


**Fig. S8. Length distribution of E. coli RNA transcripts.** Histogram of all gene transcript lengths in the *E. coli* genome (Genbank CP053602.1; **SI Appendix, Dataset S2**). Arrows show the length of pcoat′ and pMS2′.

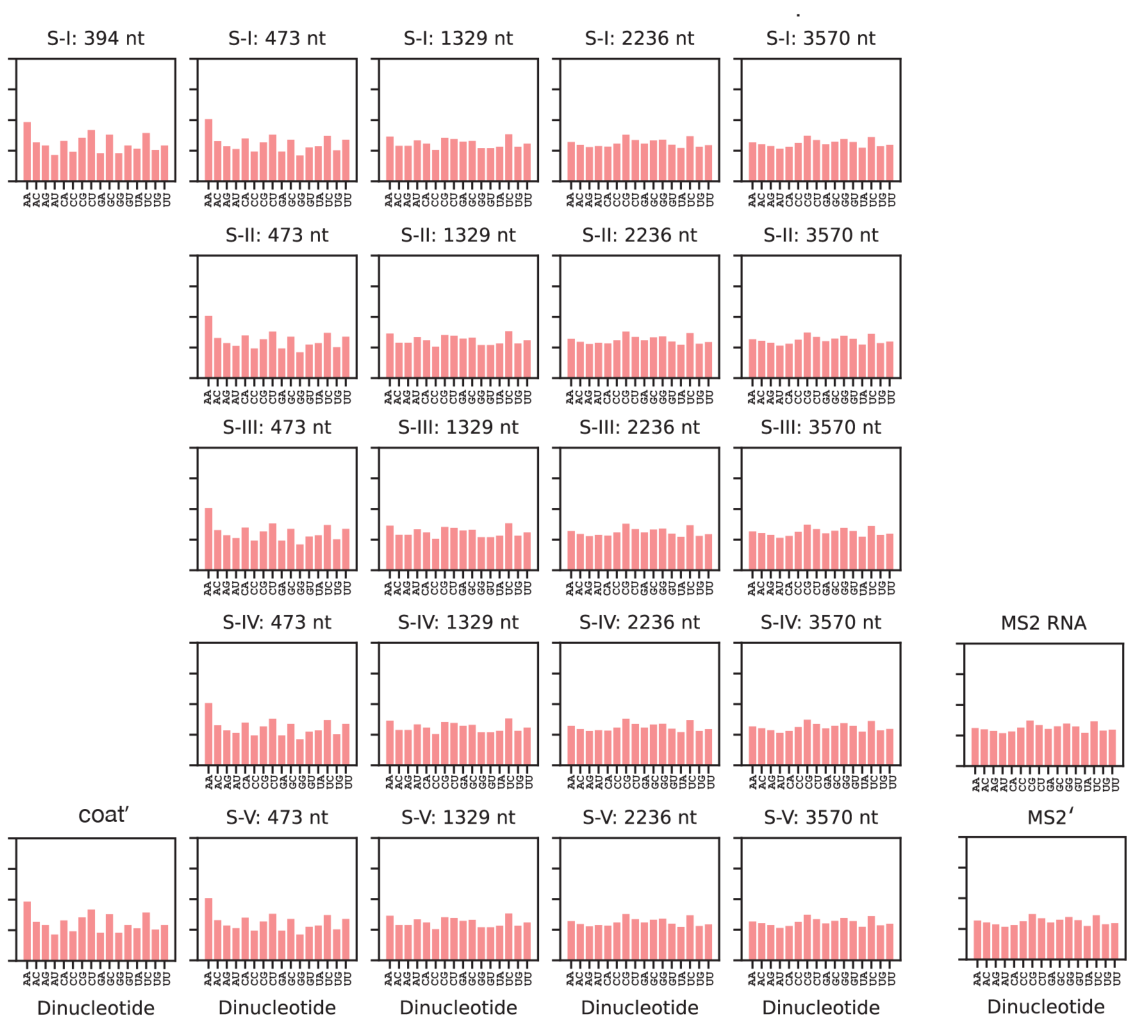


**Fig. S9. Dinucleotide frequencies of RNA transcripts.** Dinucleotide frequencies for each transcript length are conserved across shuffle types.

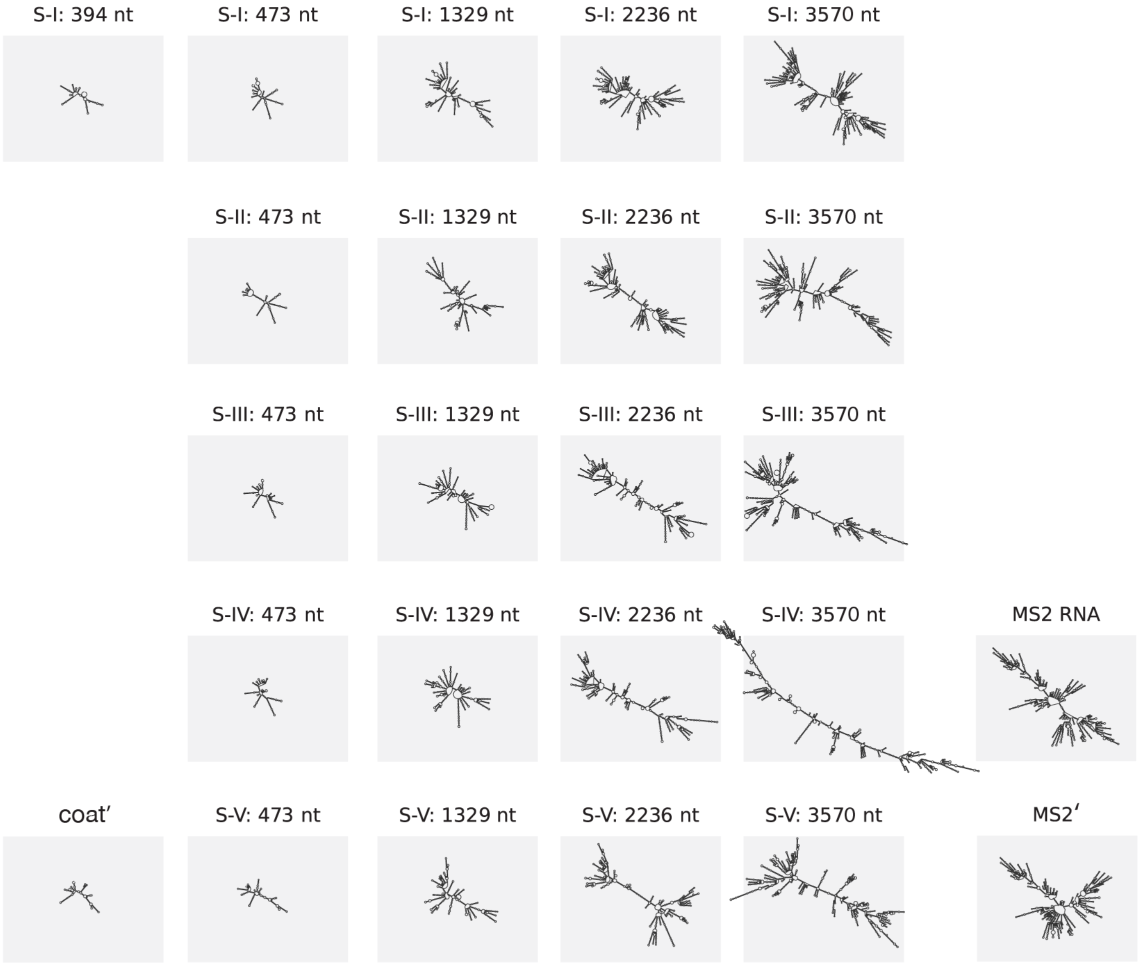


**Fig. S10. Theoretical MFE structures of RNA transcripts predicted by RNA folding models.** We used RNAfold from the ViennaRNA package (36) to predict the minimum-free-energy (MFE) secondary structure of each insert transcript. While the MFE structure does not directly reveal the physical structure of an RNA molecule, it provides some information about the intramolecular base pairing and overall extendedness of the molecule.

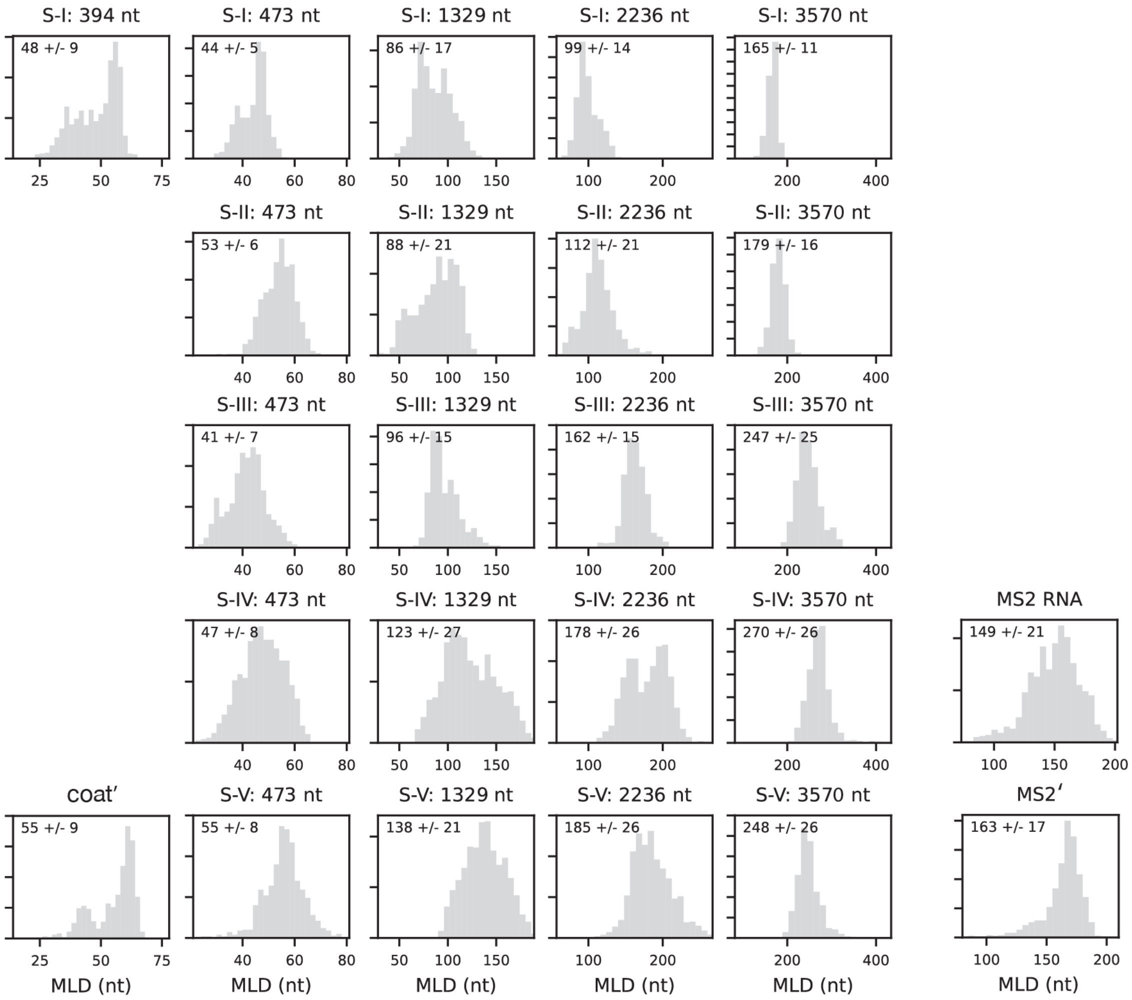


**Fig. S11. Maximum ladder distance (MLD) of RNA transcripts.** The MLD, derived from RNA secondary structure models, estimates the physical size of each RNA transcript. We computed MLD values for 1,000 suboptimal structures of each insert sequence.

| Shuffle type | Insert length | Biological replicate | Insert coverage | Vector coverage | Host coverage | Total coverage | Insert % | Vector % | Host % |
|---|---|---|---|---|---|---|---|---|---|
| pMS2' | 3574 | - | 1.28e8 | 5.68e5 | 3.12e6 | 1.32e8 | 97.2% | 0.4% | 2.4% |
| pMS2' | 3574 | duplicate | 1.38e8 | 5.15e5 | 2.89e6 | 1.42e8 | 97.6% | 0.4% | 2.0% |
| pMS2' - no pack | 3574 | - | 2.07e6 | 1.86e3 | 1.38e8 | 1.40e8 | 1.5% | 0.0% | 98.5% |
| pcoat' | 394 | - | 2.54e6 | 2.76e7 | 6.69e7 | 9.71e7 | 2.6% | 28.4% | 68.9% |
| pcoat' | 394 | duplicate | 5.31e6 | 3.53e7 | 9.47e7 | 1.35e8 | 3.9% | 26.1% | 70.0% |
| SI | 394 | - | 8.69e6 | 2.66e7 | 6.11e7 | 9.64e7 | 9.0% | 27.6% | 63.3% |
| SI | 394 | duplicate | 8.61e6 | 3.30e7 | 9.68e7 | 1.38e8 | 6.2% | 23.8% | 70.0% |
| SI | 394 | triplicate | 1.15e7 | 4.00e7 | 9.12e7 | 1.43e8 | 8.1% | 28.0% | 63.9% |
| SI | 473 | - | 2.76e7 | 2.04e7 | 4.76e7 | 9.55e7 | 28.9% | 21.3% | 49.8% |
| SI | 473 | duplicate | 3.93e7 | 2.75e7 | 2.93e7 | 9.61e7 | 40.9% | 28.6% | 30.5% |
| SII | 473 | - | 1.44e7 | 2.21e7 | 5.96e7 | 9.62e7 | 15.0% | 23.0% | 62.0% |
| SII | 473 | duplicate | 7.56e6 | 2.92e7 | 6.03e7 | 9.70e7 | 7.8% | 30.0% | 62.2% |
| SIII | 473 | - | 4.61e7 | 1.93e7 | 2.98e7 | 9.52e7 | 48.4% | 20.3% | 31.3% |
| SIII | 473 | duplicate | 2.95e7 | 2.78e7 | 3.92e7 | 9.65e7 | 30.6% | 28.8% | 40.6% |
| SIV | 473 | - | 1.18e7 | 2.43e7 | 6.10e7 | 9.71e7 | 12.1% | 25.1% | 62.8% |
| SV | 473 | - | 7.09e6 | 2.63e7 | 6.39e7 | 9.72e7 | 7.3% | 27.0% | 65.7% |
| SI | 1329 | - | 5.92e7 | 8.11e6 | 2.68e7 | 9.41e7 | 62.9% | 8.6% | 28.4% |
| SII | 1329 | - | 2.36e7 | 2.74e7 | 4.54e7 | 9.64e7 | 24.4% | 28.4% | 47.1% |
| SIII | 1329 | - | 3.87e7 | 2.08e7 | 3.76e7 | 9.70e7 | 39.9% | 21.4% | 38.7% |
| SIV | 1329 | - | 1.40e7 | 3.50e7 | 4.57e7 | 9.47e7 | 14.8% | 37.0% | 48.3% |
| SV | 1329 | - | 8.98e6 | 4.33e7 | 4.51e7 | 9.74e7 | 9.2% | 44.5% | 46.3% |
| SI | 2236 | - | 7.74e7 | 2.94e6 | 1.44e7 | 9.47e7 | 81.7% | 3.1% | 15.2% |
| SII | 2236 | - | 3.04e7 | 2.74e7 | 3.92e7 | 9.70e7 | 31.3% | 28.3% | 40.4% |
| SIII | 2236 | - | 5.32e7 | 1.48e7 | 2.80e7 | 9.60e7 | 55.4% | 15.4% | 29.2% |
| SIV | 2236 | - | 1.71e7 | 5.55e7 | 2.46e7 | 9.73e7 | 17.6% | 57.1% | 25.3% |
| SV | 2236 | - | 1.22e7 | 5.76e7 | 2.70e7 | 9.68e7 | 12.6% | 59.5% | 27.9% |
| SI | 3570 | - | 8.19e7 | 1.79e6 | 8.48e6 | 9.22e7 | 88.9% | 1.9% | 9.2% |
| SI | 3570 | duplicate | 7.94e7 | 3.34e6 | 1.22e7 | 9.50e7 | 83.6% | 3.5% | 12.8% |
| SIII | 3570 | - | 3.93e7 | 2.46e7 | 3.32e7 | 9.71e7 | 40.5% | 25.3% | 34.2% |
| SIII | 3570 | duplicate | 6.68e7 | 9.45e6 | 2.02e7 | 9.64e7 | 69.2% | 9.8% | 21.0% |
| SIV | 3570 | - | 3.02e7 | 4.19e7 | 2.49e7 | 9.70e7 | 31.1% | 43.2% | 25.7% |
| SV | 3570 | - | 2.49e7 | 4.16e7 | 3.05e7 | 9.70e7 | 25.6% | 42.9% | 31.5% |
| SV | 3570 | duplicate | 2.35e7 | 4.77e7 | 2.58e7 | 9.69e7 | 24.2% | 49.2% | 26.6% |
| SIV+3 | 3570 | - | 7.72e7 | 1.65e7 | 4.28e7 | 1.37e8 | 56.5% | 12.1% | 31.4% |
| SIV+1 | 3570 | - | 6.71e7 | 3.17e7 | 4.00e7 | 1.39e8 | 48.3% | 22.8% | 28.8% |
| SI-3 | 3570 | - | 1.29e8 | 4.36e6 | 7.66e6 | 1.41e8 | 91.5% | 3.1% | 5.4% |
| SI-1 | 3570 | - | 1.20e8 | 2.42e6 | 1.48e7 | 1.38e8 | 87.5% | 1.8% | 10.7% |
| natural MS2 | 3569 | - | 9.50e7 | - | 1.53e5 | 9.52e7 | 99.8% | - | 0.2% |

**Table S1.** Table of RNAseq coverage analysis performed in this study.

| Data collection and processing | pMS2' T=3 | pMS2' D5 | pcoat' T=3 | pcoat' D5 | pcoat' oblate |
|---|---|---|---|---|---|
| EMDB ID | EMD-48864 | EMD-48866 | EMD-48865 | EMD-48867 | EMD-48868 |
| PDB ID | 9N40 | - | 9N41 | - | - |
| # micrographs | 7,103 | 7,103 | 6,833 | 6,833 | 6,833 |
| # final particles | 728,471 | 4,976 | 313,055 | 6,932 | 19,577 |
| Symmetry imposed | I1 | D5 | I1 | D5 | C1 |
| Å Resolution ($FSC_{0.143}$) | 2.4 | 3.5 | 2.2 | 3.3 | 3.5 |

**Table S2.** Table of cryo-EM data.

| **Refinement stats (asu)** | **pMS2' T=3** | **pcoat' T=3** |
|---|---|---|
| PDB ID | 9N40 | 9N41 |
| Bonds (RMSD) | | |
| Length (Å) | 0.002 | 0.002 |
| Angles (°) | 0.450 | 0.458 |
| MolProbity score | 0.90 | 0.90 |
| Clash score | 1.56 | 1.56 |
| Rotamer outliers (%) | 0.63 | 0.31 |
| Ramachandran plot (%) | | |
| Outliers | 0.00 | 0.00 |
| Allowed | 1.06 | 0.79 |
| Favored | 98.94 | 99.21 |
| CC (volume) | 0.95 | 0.96 |

**Table S3.** Table with additional cryo-EM data.
